# Discovery and optimization of the next generation of cell active Protein Kinase Novel 3 (PKN3) inhibitors

**DOI:** 10.64898/2026.08.20.745975

**Authors:** Eleni Georgiou, Tuomo Laitinen, Antti Poso, Rosemarie Heino, Christopher R. M. Asquith

## Abstract

Protein Kinase Novel 3 (PKN3) understudied kinase with a diverse array of biological functions that are yet to be fully defined. Here, we report the design and development of a novel advanced functional chemical tool inhibitor for PKN3. A pyridyl imidazole series has been synthesized and evaluated against PKN3 *in vitro* and in cells. These efforts led to the discovery of **6e** (URS03-06), a submicromolar cell active functional inhibitor with a narrow kinome spectrum, to enable the elucidation and interrogation of PKN3 cellular biology.

**Graphic:** Shining light on Protein Kinase Novel 3 (PKN3), an understudied kinase with a wide array biological functions that are yet to be fully explored. We report on the design and development of a novel advanced functional chemical tool inhibitor for PKN3 URS03-06 (**6e**). This pyridyl imidazole series has been synthesized and evaluated against PKN3 in vitro, in cells and evaluated across the kinome. The resulting compound, URS03-06 (**6e**) is a narrow spectrum inhibitor of with an IC_50_ of 210nM in cells and a narrow kinome profile S_30_=0.02 at 1μM. The discovery of URS03-06 (**6e**) could enable the further elucidation and interrogation of PKN3 cellular biology.

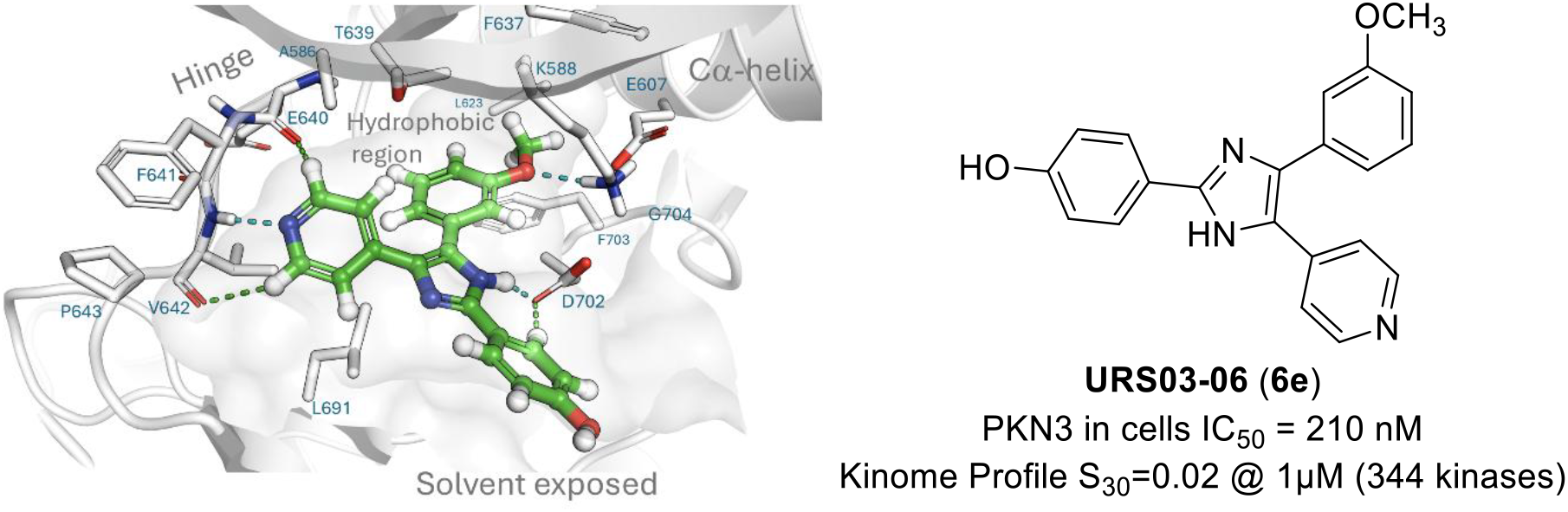

---

Protein Kinase Novel 3 (PKN3, also known as PKNβ and PRK3) is one of the three serine/threonine isozymes PKN1-3 that are part of the PKN subfamily which belong to the AGC protein kinase superfamily [1]. The PKNs isozymes share a homologous catalytic domain to Protein Kinases C (PKCs) at their C-terminal region within the same superfamily and are often referred to as PRKs (protein kinase C-related kinases) [2, 3]. PKN1-3 are characterized by an N-terminal region containing three antiparallel coiled-coil (ACC) repeats and a C2-like domain [2, 4]. The unique domains and sequence features of PKN3 confer distinct regulatory mechanisms and biological functions to PKN1 and PKN2. While the three isozymes share a similar structural organization, they have significant differences in both tissue distribution [4, 5, 6] and regulatory mechanisms, including their response to stiluli, fatty acids, phosphoinositides, and distinct effectors [4, 7, 8]. PKN1 and PKN2 are ubiquitously expressed across a wide range of mammalian tissues. In contrast, PKN3 mRNA expression is restricted to a limited number of normal tissues, including skeletal muscle, heart, and liver, where it is generally detected at low levels, as well as in endothelial cells [2, 6]. Notably, PKN3 is highly expressed in several human cancer cell lines, suggesting a distinct role in tumor biology [3, 9].

The PKN family has also been found to be the first effector kinases of the Rho family of the small GTPases, which are important in regulating cytoskeletal dynamics and cell mobility including migration and adhesion [10]. However, PKN1-3 differ in responsiveness to Rho of the GTPases, phospholipids, fatty acids and arachidonic acid [4, 7, 8]. The physiological role of PKN3 was investigated by Mukai *et al*. using PKN3 knockout (KO) mice and demonstrated that PKN3 was not essential for normal development, growth to adulthood, or physiological vascular development [7]. However, embryonic fibroblasts derived from PKN3 KO mice exhibited markedly reduced migratory activity compared with wild-type (WT) cells in Boyden chamber-based transwell migration assays; a defect that was consistently observed across multiple growth factor-induced migration models. These findings indicate that PKN3 contributes to cell motility by regulating actin cytoskeleton organization in primary fibroblasts. Although PKN3 also supports angiogenesis, its deletion did not inhibit tumor angiogenesis, suggesting that compensatory mechanisms or alternative signaling pathways can sustain tumor-associated blood vessel formation in the absence of PKN3 [7].

PKN3 is although increasingly recognized as a key contributor to cancer progression, particularly in processes associated with tumor invasion and metastasis [3, 9]. It participates in signaling networks through interactions with RhoC [11] and p130Cas, both of which regulate cell migration, proliferation, and invasiveness [12]. As a downstream effector of the phosphoinositide 3-kinase (PI3K) pathway, PKN3 is activated following PI3K signaling. Under physiological conditions, PI3K activity is tightly controlled by the tumor suppressor PTEN [5]. However, loss or inactivation of PTEN frequently occurrs in human cancer, resulting in sustained PI3K signaling and increased PKN3 expression, thereby promoting metastatic behavior [12]. Experimental evidence supports a functional role for PKN3 in multiple cancer types. Elevated exogeneous PKN3 expression enhances the aggressive characteristics of breast cancer cells *in vitro* [11], while inhibition of PKN3 suppresses the growth of PI3K-dependent prostate and breast tumor xenografts [5, 11, 13]. Since, PKN3 functions downstream of PI3K, selectively targeting this kinase may offer a strategy to inhibit oncogenic PI3K signaling while potentially reducing the adverse effects associated with direct PI3K inhibition [5]. In addition to its role in cancer, PKN3 has been implicated in the regulation of neovascularization, suggesting that it may also represent a therapeutic target for angiogenesis-related disorders, including age-related macular degeneration [7].

PKN3 may also represent a potential therapeutic target for a range of diseases, including leukemia [13-15], arthritis, age-related macular degeneration [7], rheumatoid arthritis, and osteoporosis [16]. The PI3K pathway has been reported to be overactivated in several types of leukemia, with elevated PKN3 expression observed in human T-cell acute lymphoblastic leukemia (T-ALL). Notably, PKN3 deletion delayed T-ALL progression without affecting normal hematopoiesis [13]. PKN3 has also been identified as a potential regulator of neovascularization, suggesting a possible role in vascular diseases such as arthritis and age-related macular degeneration [16]. Furthermore, PKN3 functions downstream of the Wnt5a– Ror2–Rho signaling pathway and contributes to bone resorption, supporting its potential as a therapeutic target for bone-related disorders, including rheumatoid arthritis and osteoporosis [16].

A significant advancement towards accessing PKN3’s translational potential in precision oncology was the development of a liposomal siRNA formulation, known as Atu027 [17, 18, 19]. Atu027 has been shown to silence PKN3 *in vivo* preventing liver and lung metastasis, as well as inhibiting prostate and pancreatic cancer growth. Atu027 had reached clinical trials in advanced solid tumors (NCT00938474) [20] and advanced pancreatic cancer (NCT01808638) [21], with the later showing that Atu027 in combination with gemcitabine (antimetabolite) have a favorable safety profile and enhanced clinical benefits [22]. Despite the notable progress, challenges remain with Atu027, including dose limiting toxicity, along with concerns around potential enzymatic instability, off-target effects and innate immune responses [23-25].

Alternative therapeutic strategy targeting PKN3 to RNA interference technologies is the development of small molecule kinase inhibitors. The literature landscape on such chemical tool inhibitors is currently limited. This is in part due to the lack of availability of assays and has meant PKN3 was not included in major kinome screening panels in the past, restricting data availability [26, 27, 28, 29, 30, 31, 32]. However, several PKN3 inhibitors were identified, through a number of screening approaches including focused and some broader panel screening (Figure 1). This resulted in the identification of PKN3 off-target activities on existing kinase inhibitors. Specifically, fasudil (K_i_ = 110 nM) [33], H-8 (K_i_ = 10 nM) [33], SB202190 (K_i_ = 4.0 nM) and PP1 (K_i_ = 1.3 nM) [33] have been all found to inhibit PKN3, albeit with limited wider kinome selectivity screening. In 2019, an unbiased chemoproteomic screen led to the discovery of the covalent PKN3 inhibitor JZ128, which exhibited moderate potency (IC_50_ = 120 nM) and a relatively narrow kinome inhibition spectrum [34]. More recently a large-scale kinobead-based chemoproteomic profiling (KinoBead) led to the identification of three additional, reversible PKN3 inhibitors, GSK902056A (*K*_d_^app^ of 1 nM), GSK949675A (*K*d^app^ of 2.5 nM) and SB-476429-A (*K*d^app^ of 36 nM) although the latter exhibited lower kinome selectivity (**Figure 1,a**) [35]. Despite this progress, there remains a need for more potent and selective PKN3 inhibitors. Achieving this will require systematic efforts in PKN3-targeted drug development, including comprehensive structure-activity relationship (SAR) studies and kinome-wide selectivity profiling. We have previously disclosed a deep annotation of a library of 4-anilinoqun(az)olines leading to several narrow-spectrum inhibitors [36]. Compound UNC-CA94 found to potently inhibit PKN3 in a biochemical assay (IC_50_ = 14 nM) and with micromolar activity in cells (IC_50_ = 1.3 μM).

**Figure 1.**
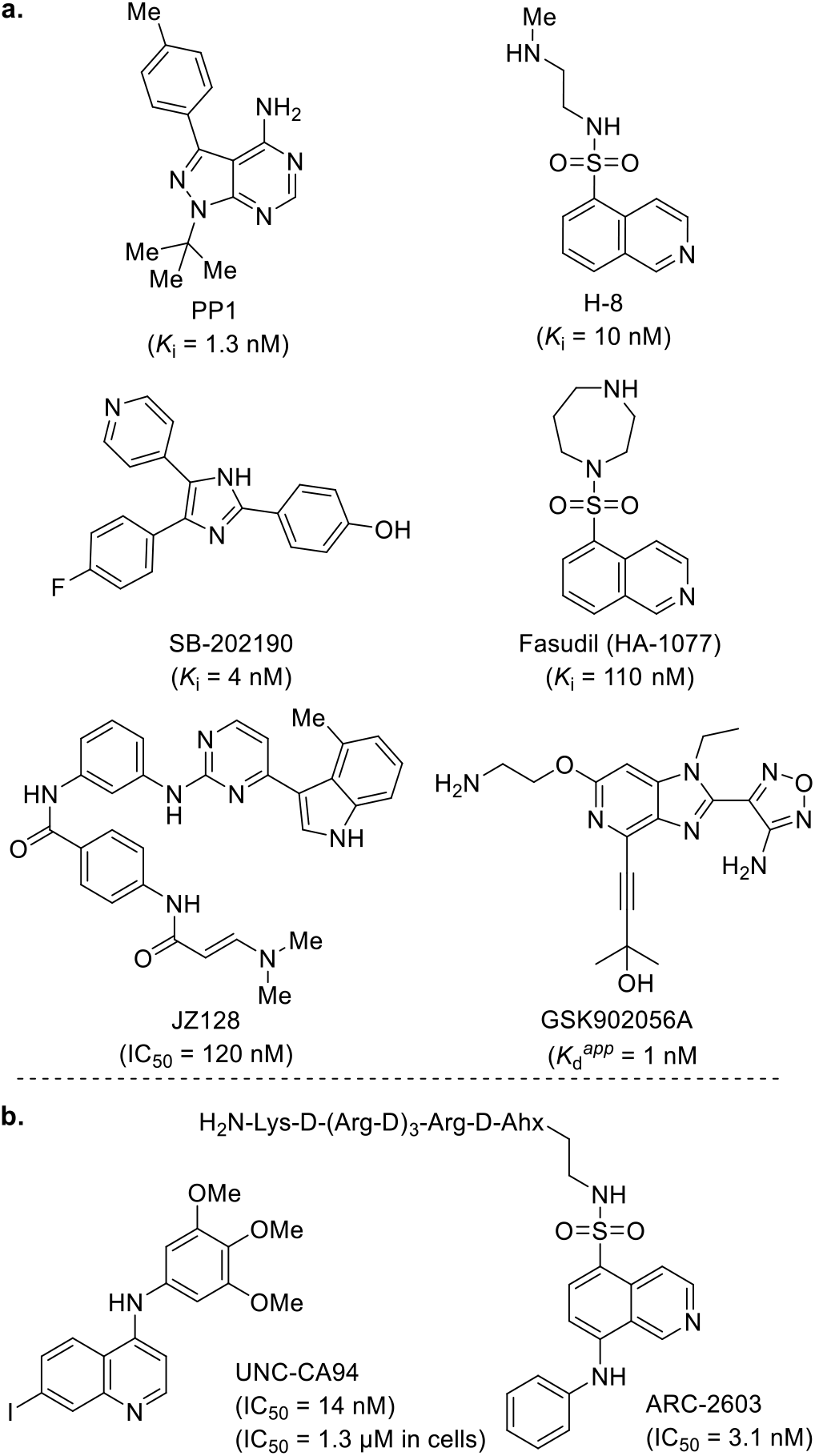
Previously reported compounds active against PKN3: a. resulted from repurposing/kinome-wide screening; b. resulted from SAR studies

More recently, Smorodina *et al*. reported a bisubstrate inhibitor, ARC-2603, exhibiting potent biochemical inhibition (IC_50_ = 3.1 nM) against PKN3 [37]. Nevertheless, ARC-2603’s in-cell target engagement was not reported. The selectivity panel, conducted across 397 protein kinases, showed that in addition to inhibiting PKN3, ARC-2603 also strongly inhibits the tyrosine kinase DDR1 as well as the PKN1 subfamily member (**Figure 1,b**). Hence, to date there has not been a cell-active chemical probe quality compound disclosed in the literature [38].

## Results

We aimed to identify a tractable chemical starting point by evaluating previously reported compounds with activity against PKN3 (**Figure 1**). The identification of a potent compound exhibiting high selectivity for PKN3 within the PKN subfamily was considered critical. SB-202190 has been reported to display exceptional potency (K_i_ = 4.0 nM) along with pronounced selectivity for PKN3 over PKN1 and PKN2. Notably, this compound, originally developed as a p38 MAPK inhibitor, is characterized by a pyridyl imidazole scaffold. Having this structural feature in hand, we screened additional pyridyl imidazole derivatives from the same literature SB series (**Figure 2**) for their activity against PKN3. Selected examples from the SB series and staurosporine were screened in a PKN3 enzyme assay [29], which enabled us to quickly map the PKN3 inhibition landscape.

**Figure 2.**
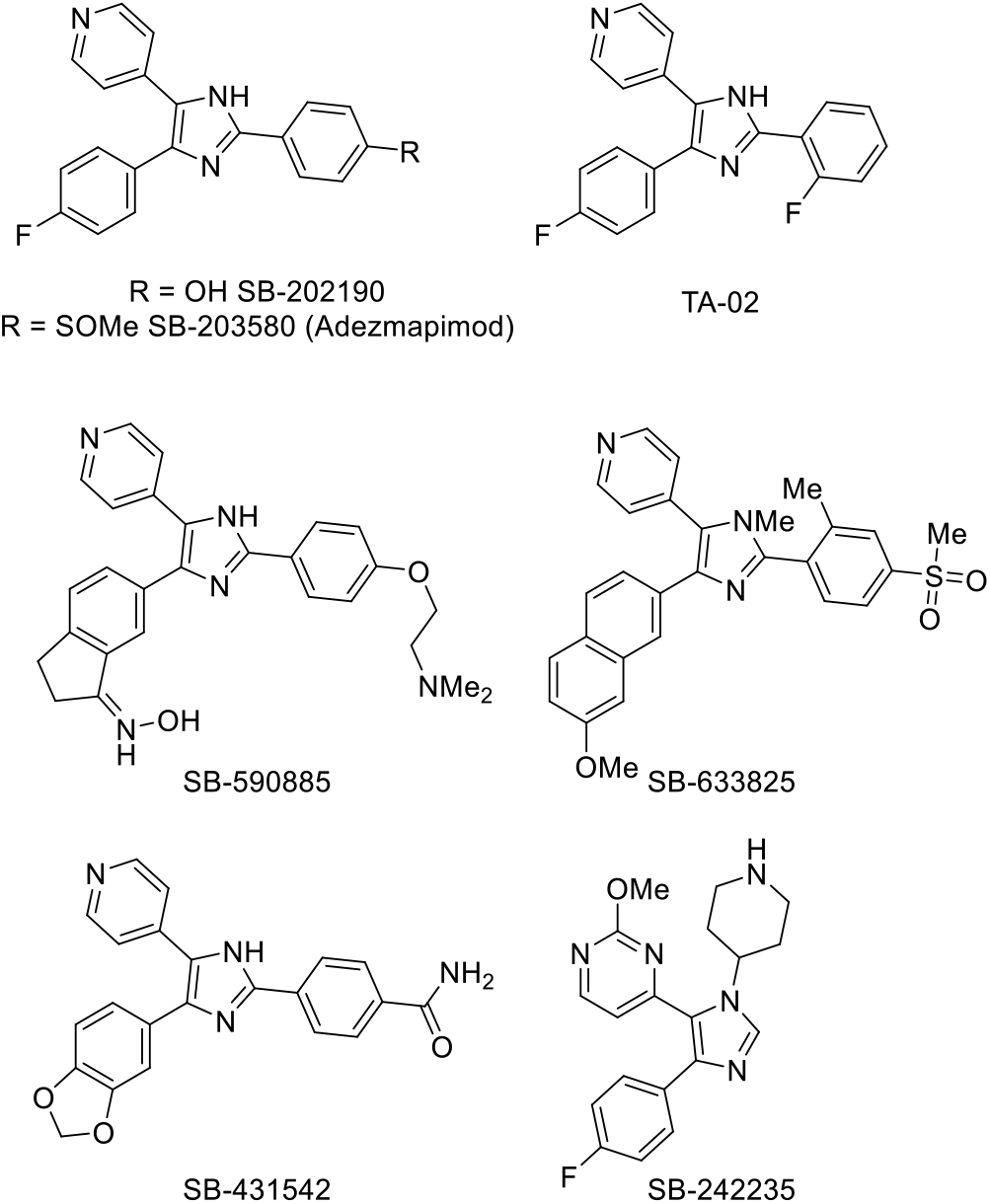
Structures of imidazoles tested that were previously reported as inhibitors against other kinases.

The compounds were tested in PKN3 enzyme assay in 10-point dose dependent format to determine the IC_50_ values. The maximum concentration used was 20 μM with an ATP concentration of 10 μM. The imidazoles had a range of activity, most exhibited low single digit micromolar inhibition against PKN3 (**Table 1**). The starting point compound, SB-202190 had good potency against PKN3 (IC_50_ = 0.34 μM). Surprisingly, the switch from the phenolic alcohol of SB-202190 to the methyl sulfoxide of SB-303580 (Adezmapimod) resulted in a >20-fold decrease in inhibition on PKN3, this despite literature binding assays suggesting they were equipotent. We rationalized that the phenolic alcohol was able to more favorably interact with the hydrogen bonding water network [39, 40].

**Table 1.** Screening of literature imidazole scaffolds against PKN3.

| Compound | PKN3 IC<sub>50</sub> (μM)<sup>a</sup> |
| --- | --- |
| **SB 202190** | 0.34 |
| **Adezmapimod** | 8.0 |
| **TA-02** | 1.8 |
| **SB-633825** | 3.0 |
| **SB-431542** | >20 |
| **SB 242235** | 10.3 |
| **SB-590885** | 4.0 |
| **Staurosporine** | 0.0037 |

<sup>a</sup>IC<sub>50</sub> values in an enzyme assay (n=1) [29].

The *ortho*-fluoro analog of SB-202190 recovered some activity over SB-203580, with 4-fold improvement, but still a 5-fold decrease on SB-202190. SB-633825 with several modifications including a hydroxylamine showed no improvement over SB-203580. SB-431542, which is an analog of SB-303580 with several modifications, showed no activity at the concentrations tested. SB-242235 was also a structural analog of SB-431542 and was able to retain the same level of inhibition against PKN3. SB-590885 while still an imidazole-based compound, was structurally quite distinct with a pyrimidine type hinge binder and different structural arrangement with the 2-substitution moved to the NH of the imidazole in the 1-position. SB-590885 demonstrated some flexibility within the chemotype with an IC_50_ = 4.0 μM. These results indicated that 2,4,5-substituted analogs, with 2-,4-aryl substitution pattern, constitute a more promising structural class for further optimization. Having these results in hand, we proceeded to design a series of analogs to establish preliminary structure–activity relationships (SAR) and to identify suitable synthetic routes for their preparation. Emphasis was placed on evaluating the influence of small substituents on the 4-aryl ring (zone A) in the hydrophobic pocket of the ATP binding site and the substituents on the 2-aryl ring in the solvent exposed region (zone B) (**Figure 3**)

**Figure 3.**
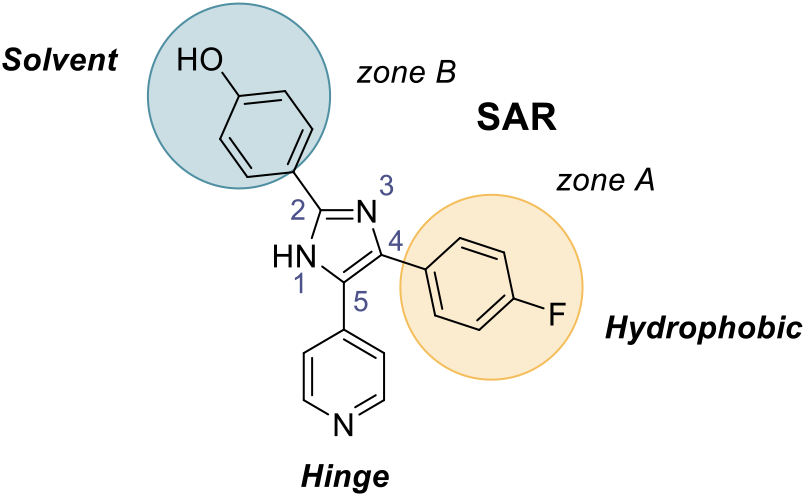
Design rationale to assess the imidazole chemotype.

Two principal synthetic routes were employed to access the kinase inhibitor cores in compounds **6a-o, 14a-f** and **16**, (**Scheme 1** and **Scheme 2**). The strategies are largely based on previously reported methods for the synthesis of pyridyl imidazole hybrid scaffolds [41, 42, 43]. In general, 4-substituted analogs were prepared *via* **Scheme 1**, which involves oxidation of **1** followed by condensation with aryl aldehydes **2** to construct the central imidazole ring in **3**. Subsequent bromination of the imidazole enabled installation of the aryl substituent at the C4 position through Suzuki–Miyaura cross-coupling reactions with boronic acids **5a-m** to furnish compounds **6a-m**. Interestingly, when boronic acid **5i** subjected to the reaction conditions, the hydrated byproduct **6i’** was formed along with the expected product **6i**. In cases where 2-ether-containing analogs were targeted, these were obtained *via* post-functionalization of intermediate **6a. Scheme 2**, based on modified procedures from the original synthesis of SB-202190, was utilized to access 2-amine-containing analogs. In this approach, a Weinreb reaction between an aryl Weinreb amide **7** and 4-methylpyridine **8** afforded the corresponding ketone **9**. The resulting α-keto oxime **10**, generated under oxidative nitration conditions, was subsequently condensed with an aryl aldehyde **11** in the presence of ammonium acetate to form the central imidazole ring in **12**. Final functionalization *via* Buchwald–Hartwig cross-coupling with secondary amines **13a-f** furnished the desired analogs (**14a-f**). The final analog **16**, was synthesized *via* a condensation of oxime **10** with commercially available aldehyde **15**, followed by a dihydroxylation step.

The synthesized compounds were then tested using the same PKN3 enzyme assay in 10-point dose dependent format to determine the IC_50_ values. First, we tested the influence of small functional group substituents (*ortho*-, *meta*-, *para*-) on the aryl ring (zone A) at the 4-position of the imidazole ring (**Table 2**). In the first instance, we followed up on the promising hit compound SB-202190 which when resynthesized **6a**, has showed an IC_50_ = 0.41 μM against PKN3. Transferring the fluoro group from the *para*-to the *meta*-position, yielding compound **6b**, resulted in a 2-fold decrease in potency against PKN3. In contrast, the *ortho*-position fluoro **6c**, had a 10-fold potency boost with respect to **6a**. Introduction of a methoxy group at the *para*-position **6d** was equipotent to **6a**. The *meta*-methoxy analogue **6e** was 4-fold less potent, while the corresponding *ortho*-substituted analogue **6f** showed an 11-fold improvement against PKN3 compared to **6a**.

**Table 2.** Results of ‘Zone A’ optimization of SB-202190 (**6a**).

| Compound | R<sup>1</sup> | R<sup>2</sup> | R<sup>3</sup> | PKN3<sup>a</sup> |
| --- | --- | --- | --- | --- |
|  |  |  |  | IC<sub>50</sub> (μM) |
| **6a** | F | H | H | 0.41 |
| **6b** | H | F | H | 0.94 |
| **6c** | H | H | F | 0.039 |
| **6d** | OCH<sub>3</sub> | H | H | 0.42 |
| **6e** | H | OCH<sub>3</sub> | H | 1.5 |
| **6f** | H | H | OCH<sub>3</sub> | 0.036 |
| **6g** | CN | H | H | 3.3 |
| **6h** | H | CN | H | 14.4 |
| **6i** | H | H | CN | 1.7 |
| **6i'** | H | H | CONH<sub>2</sub> | >20 |
| **6j** | COCH<sub>3</sub> | H | H | 3.6 |
| **6k** | CONHCH<sub>3</sub> | H | H | >20 |
| **6l** | NHCOCH<sub>3</sub> | H | H | 4.1 |
| **6m** | SO<sub>2</sub>CH<sub>3</sub> | H | H | 10.3 |

<sup>a</sup>IC<sub>50</sub> values in an enzyme assay (n=1) [29].

Switching to a cyano group substitution, afforded a 8-fold less potent analog **6g**, compared to the homologues *para*-substituted analogs **6a** and **6d**. The relocation of the cyano group at the meta-position **6h**, resulted in an even more dramatic decrease in potency against PKN3 (IC_50_ = 14.4 μM). The *ortho*-cyano analog **6i**, exhibited a 4-fold drop in PKN3 potency compared to **6a**. The related *ortho*-primary amide **6i’** showed no activity at the concentrations tested (IC_50_ = >20 μM). Additional *para*-substituted analogs including the acetyl **6j**, methyl carboxamide **6k**, acetamido **6l** and methyl sulfone **6m** all demonstrated weaker activity against PKN3. Compound **6j** and **6l** had low signal digit micromolar activities where, **6k** and **6m** were weaker (IC_50_ = >10 μM).We continue our SAR studies with analogs bearing ether chain and amine substitutions at the C2-aryl moiety (zone B) to probe the solvent front of the ATP pocket (**Table 3**). The alkylated phenol analogs **6n** and **6o** demonstrated weaker activity (IC_50_ = >10 μM) against PKN3. Switching to a methyl piperazine **14a**, improved inhibition was still but was still 8-fold weaker than **6a**. In contrast, the morpholine analog **14b**, was inactive at the concentrations tested (IC_50_ = >20 μM). Introduction flexibility with an acyclic amine, tetramethylethane-1,2-diamine-ethane **14c**, was also detrimental to activity against PKN3 (IC_50_ = >20 μM). Some improvements were observed with the introduction of a chiral methyl substituted piperazines analogs **14d** and **14e** resulted in a 10- and 13-fold decrease over **6a** respectively. Finally, the removal of the N-methyl of the piperazine group on **14a** to afford **14f** led to an inferior result against PKN3 as did removal of the ring to afford the diethylamine analog **16**.

**Table 3.** Results of ‘Zone B’ optimization of SB-202190 (**6a**).

| Compound | R | PKN3<sup>a</sup> |
| --- | --- | --- |
|  |  | IC<sub>50</sub> (μM) |
| **6n** | A | 15.9 |
| **6o** | B | >20 |
| **14a** | C | 3.3 |
| **14b** | D | >20 |
| **14c** | E | >20 |
| **14d** | F | 4.3 |
| **14e** | G | 5.1 |
| **14f** | H | 4.6 |
| **16** | I | 5.5 |

<sup>a</sup>IC<sub>50</sub> values in an enzyme assay (n=1) [29].

Having identified the most potent inhibitors (**6a-f**) in the enzyme assay, we sought to evaluate their in-cell target engagement using the NanoBRET assay using **6k** as negative control [38]. HEK293 cells were transiently transfected with PKN3-NanoLuc Fusion vector. The transfected cells were then treated in duplicate with a 10-point dose dependent format to determine the IC_50_ values. The maximum concentration used was 20 μM, the K-10 tracer had a concentration of 10 μM [40, 44, 45]. The IC_50_ values from the NanoBRET assay while consistent with the enzyme data, did not form an exact linear relationship; this is likely due to the differences in cell penetrance and the assay formats (**Table 4**). However, five out of the seven compounds exhibited submicromolar in-cell activity against PKN3. Compound **6e** (URS03-06) showed the highest potency with IC_50_ of 0.210 μM, making it the most potent PKN3 cellular inhibitors of PKN3 in the literature to date. URS03-06 (**6e**) was also nearly 14-fold more potent than the literature compound Adezmapimod against PKN3 and equipotent with the reference board spectrum kinase inhibitor CEP-701.

**Scheme 1.**
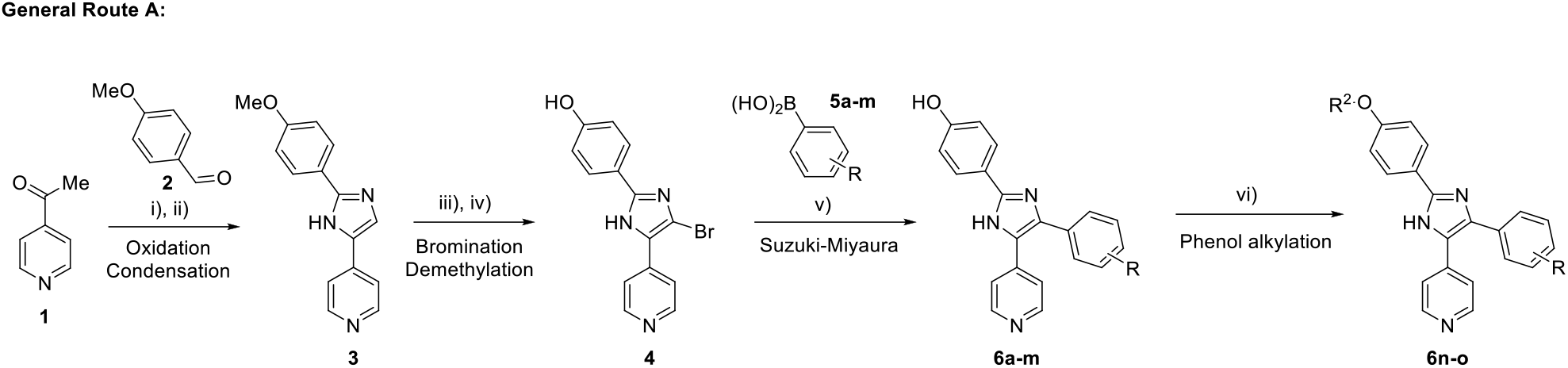
General synthetic routes to analogues (**6a-o**)-*reagents and conditions* i) aq. HBr, DMSO, 60 °C, 19 h; ii) NH_4_OAc MeOH; iii) Br_2_, Pyridine, CH_2_Cl_2_, 0 °C, 1 h; iv) 1M BBr_3_ (CH_2_Cl_2_) CH_2_Cl_2_, 0 °C to RT, 16 h; v) K_2_CO_3_, PdCl_2_(PPh_3_)_2_, DME:H_2_O, microwave 150 °C, 30-35 min; vi) 1,2-epoxy-2-methylpropane, NaHCO_3_, DMF or 1-bromo-2-methoxyethane, K_2_CO_3_, DMF.

**Scheme 2.**
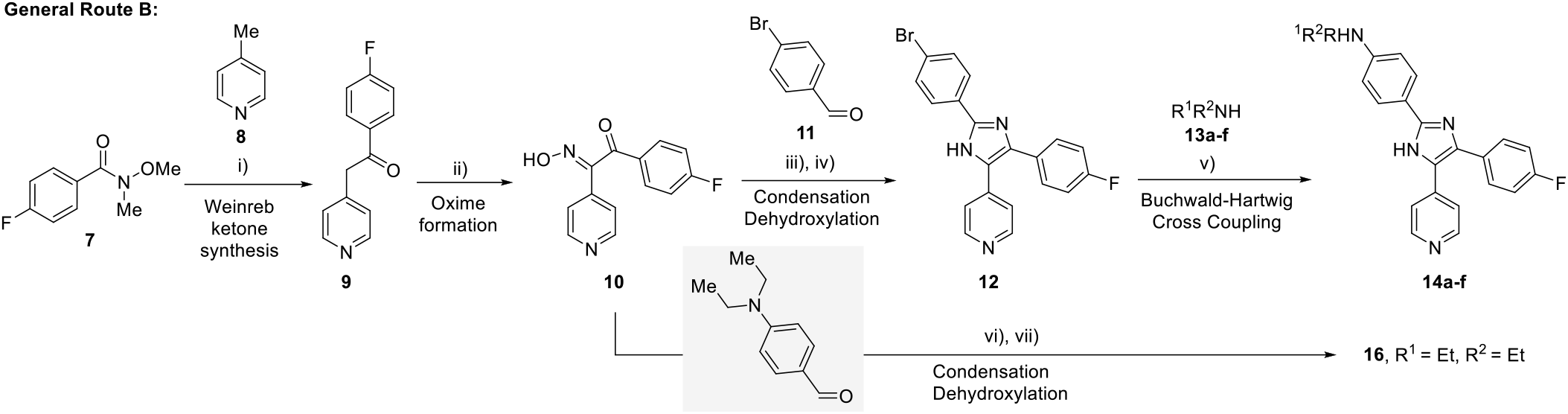
General synthetic routes to analogues (**14a-f & 16**) - *reagents and conditions* i) LDA, dry THF; ii) NaNO_2_ AcOH, H_2_O; iii) NH_4_OAC, AcOH; iv) P(OEt)_3_, DMF; v) (a) DavePhos, Pd_2_db_3_, NaOtBu, t-BuOH, 1,4-dioxane, (b) DavePhos,Pd_2_db_3_, NaOtBu, 1,4-dioxane, (c) RuPhos, Pd_2_db_3_, NaOtBu, DMA; vi) NH_4_OAc, toluene:AcOH, 110 °C, 16 h; vii) P(OEt)_3_, DMF.

**Table 4.** PKN3 NanoBRET results of the most potent inhibitors.

| Cmpd | R<sup>1</sup> | R<sup>2</sup> | R<sup>3</sup> | PKN3<sup>a</sup> |
| --- | --- | --- | --- | --- |
|  |  |  |  | IC<sub>50</sub> (μM) |
| **6a** | F | H | H | 0.476 |
| **6b** | H | F | H | 0.820 |
| **6c** | H | H | F | 0.789 |
| **6d** | OCH<sub>3</sub> | H | H | 0.427 |
| **6e** | H | OCH<sub>3</sub> | H | 0.210 |
| **6f** | H | H | OCH<sub>3</sub> | 1.79 |
| **6k** | CONHCH<sub>3</sub> | H | H | >20 |
| **Adezmapimod** | - | - | - | 2.93 |
| **CEP-701** | - | - | - | 0.198 |

<sup>a</sup>IC<sub>50</sub> values generated in NanoBRET assay (n=2).

Since the protein crystal structure of PKN3 is not available, we looked for an alternative method to model PKN3 as previously described [40]. Briefly, we compared the sequence similarity of PKN3 to the closely related sub-family kinase members PKN1 and PKN2 and with this information built a homology model. These proteins are closely related within the kinome phylogenetic tree, with a high sequence homology (PKN3 vs PKN1 66% and PKN3 vs PKN2 57%) [46]. The experimental x-ray structures of the PKN1 and PKN2 kinase domains including their overall folding have a high degree of similarity, increasing confidence in a homology model approach as developed previously [40]. The experimental structures of PKN1 also provide insights into the apo-structure, in addition to the binding mode of several co-crystallized [47].

We performed molecular dynamics simulations (3x 500 ns) to examine the binding modes of two key PKN3 inhibitors **6a** and **6e**, and the results aligned with our expectations (**Figure 4** and **5**). The pyridyl imidazole scaffold adopted a similar binding conformation observed in several other crystal structures including P38aplha (PDB: 3ZS5 and 1A9U) [48, 49]. The two key compounds **6a** and **6e** bind to the hinge region *via* H-bond interactions on the amide of Val642, similar to UNC-CA94 [40]. However, the central imidazole core and orientation of **6a** and **6e** allows for an integrated water network interaction mediated by ASP646 and ASP702, that links round to the pendant phenolic alcohol. This network interaction is complimented by mediated tautomerization of the imidazole, which gives rise to the ability for the phenolic alcohol to interact on either side of the backbone of the solvent exposed region, something not possible in the case of SB203580 (**Figure 6**). The mixed interactions of **6a** and **6e** with the catalytic lysine (Lys588) only form a small competent of the overall binding.

**Figure 4.**
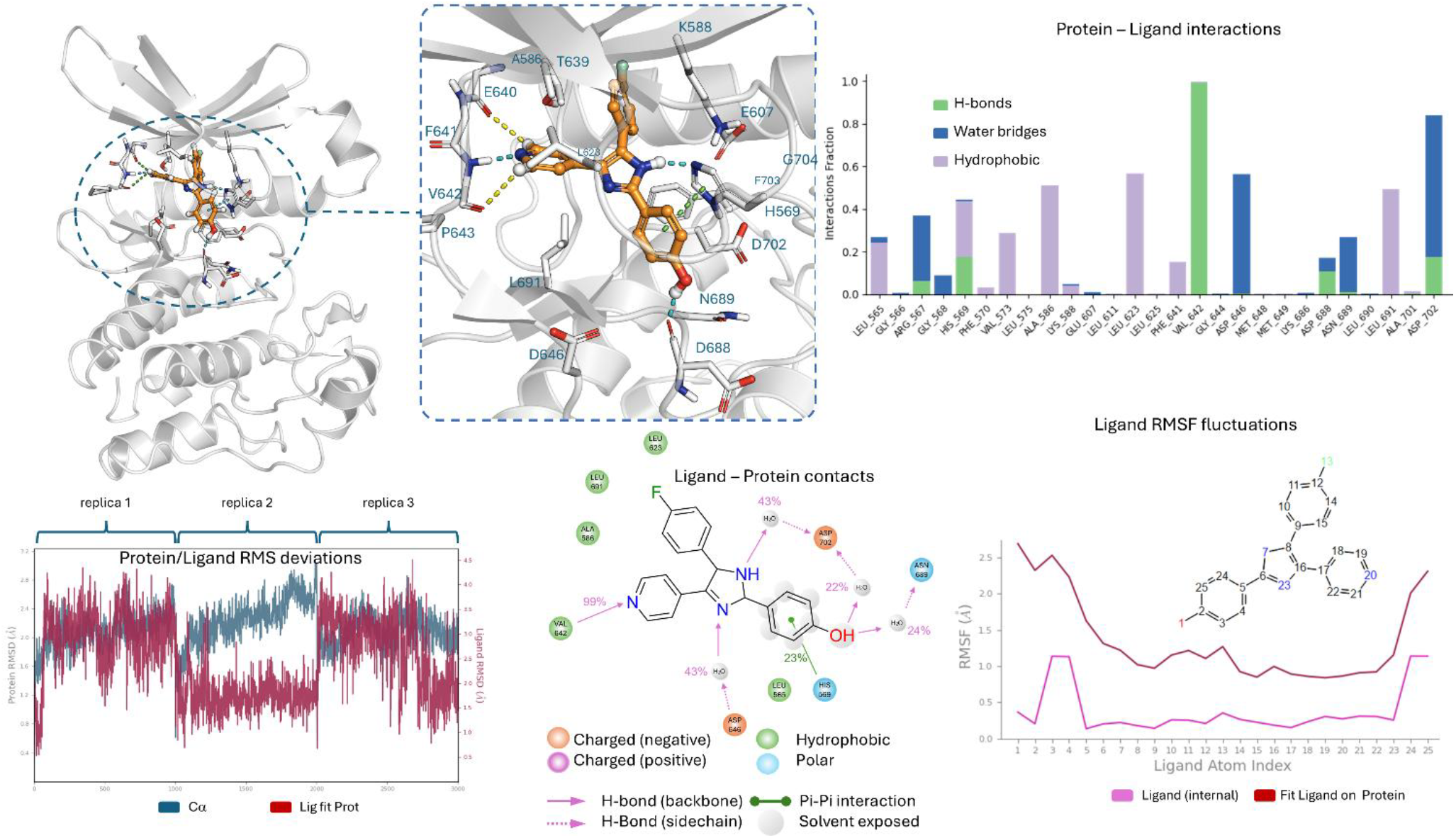
MD simulation results of **6a** with PKN3: top left: Putative binding mode of **6a** in the kinase ATP-binding pocket of PKN3 based on MD simulation; top right: Summary of main protein-ligand interactions (> 10% frequency) in the MD simulations of **6a** with PKN3; bottom left: protein/ligand interaction RMSD; middle bottom: Aggregate of protein-ligand interactions (residues with >10%) in the simulations of 6a with PKN3; top right: Ligand RMSF fluctuations in PKN3 pocket. Source data are provided in Zenodo file - https://doi.org/10.5281/zenodo.21934463.

**Figure 5.**
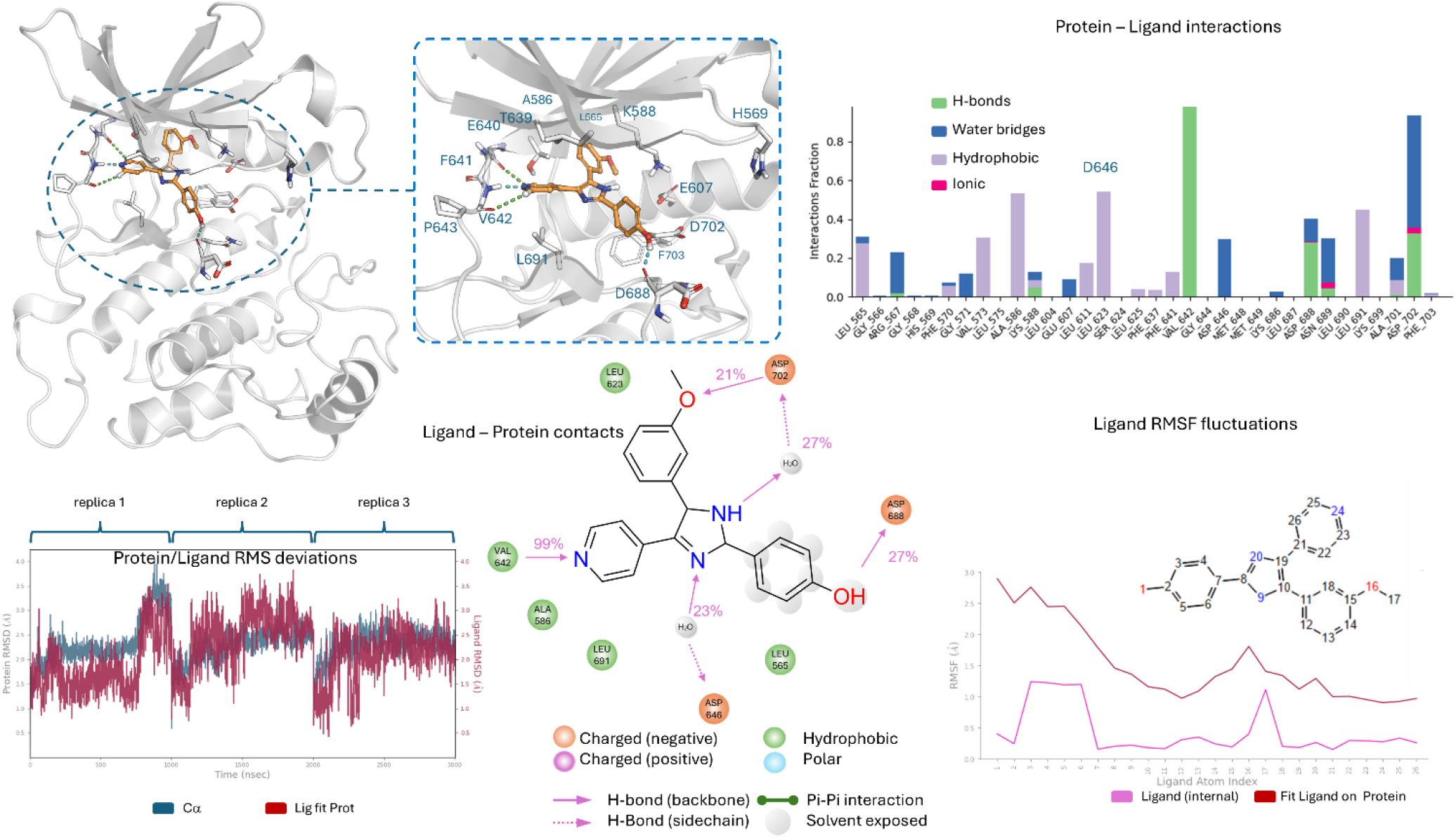
MD simulation results of **6e** with PKN3: top left: Putative binding mode of **6e** in the kinase ATP-binding pocket of PKN3 based on MD simulation; top right: Summary of main protein-ligand interactions (> 10% frequency) in the MD simulations of **6e** with PKN3; bottom left: protein/ligand interaction RMSD; middle bottom: Aggregate of protein-ligand interactions (residues with >10%) in the simulations of 6e with PKN3; top right: Ligand RMSF fluctuations in PKN3 pocket. Source data are provided in Zenodo file - https://doi.org/10.5281/zenodo.21934463.

**Figure 6.**
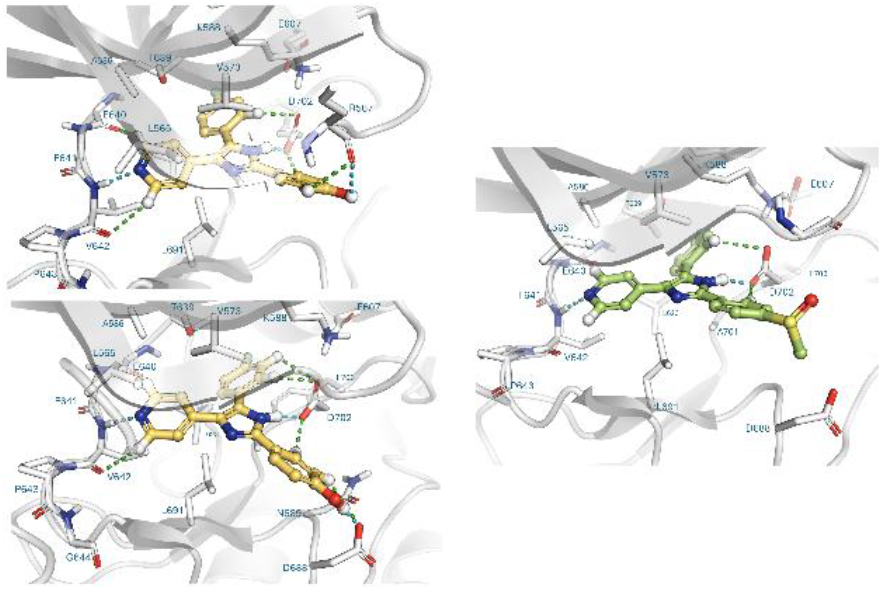
Modelling of **6a** and **6e** (top left and bottom left respectively) vs Adezmapimod (right) all docked in PKN3 highlighting the solvent exposed region interactions.

We then assessed the kinome profile of **6e** using an enzyme assay panel of 350 kinases, at a concentration of 1 μM (see supporting information, **Table S1**) [29]. The compound **6e** showed a relatively narrow spectrum of kinase inhibition and was consistent with the previous kinome profiles on related inhibitors with this scaffold (**Figure 7**). Inhibition was detected for 10 kinases above 50%, this included p38beta, RIPK2, p38alpha, NLK, PKN3, GSG2, JNK3, BRK, JNK2 and BRAF and this reduced to 7 kinases above 70%. This result demonstrated that URS03-06 (**6e**) had a narrow spectrum kinome profile.

**Figure 7.**
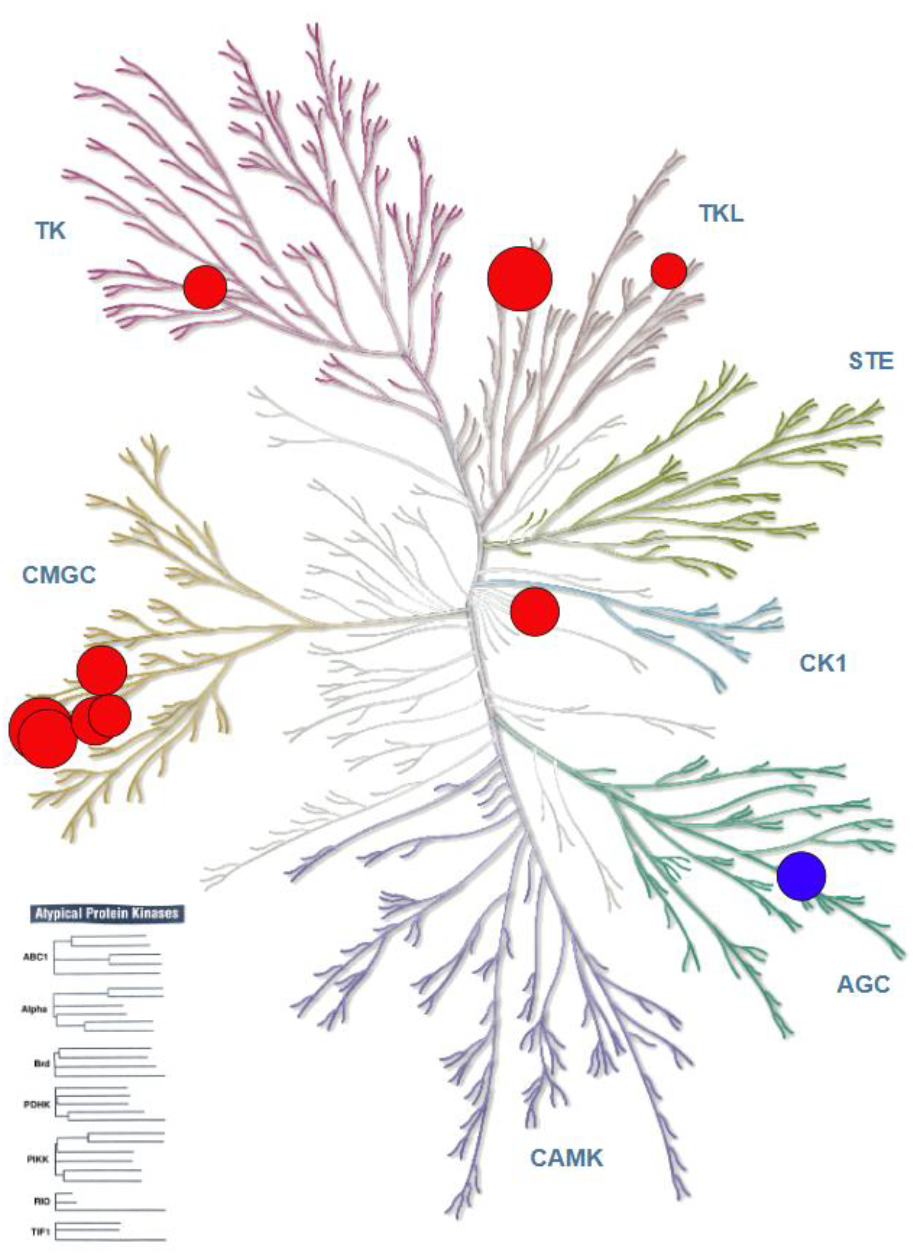
Kinome profile of **6e** against 344 kinases in a RBC enzyme assay, off-targets (>50% inhibition at 1 μM) highlighted in red with PKN3 highlighted in blue (see Table S1).

## Discussion

Protein kinases are tractable drug targets with over 100 inhibitors targeting the ATP binding site been approved by the FDA mainly for oncology but also for an increasing number of other indications [50, 51]. The effectiveness of these compounds as drugs also limits them as probes for biology. Most of these drug’s binding to more than one kinase potentially shutting down multiple parallel signaling pathways. This has limited the therapeutic use of these drugs primarily oncology indications. Inhibitors with improved potency and selectivity profiles will likely be needed to treat kinase indications beyond oncology and for more precision/personalized targeted therapy in oncology [52]. Developing potent and selective high-quality chemical probes is an excellent proof of concept to achieving these aims and derisking the therapeutic pathway.

In the case of PKN3, where experimental evidence supports a functional role for PKN3 in multiple cancer types. We started with a relatively potent literature hit compound SB-202190 [33]. Through a series of rationale design and molecular modelling we defined at series of structure-activity relationships. We were able to define the narrow cavity within the hydrophobic region of the binding site, where *ortho*-cyano analog **6i** highlighted some conformational limitations and that going beyond small substitutions were not tolerated. The pyridine-based hinge binder worked well. While we took advantage of the central water network, our extensions into the solvent exposed region were less successful. The phenolic alcohol proved to be versatile at forming a number of solvent front interactions. These interactions combined with more effective orientation in the ATP binding domain allowed us to boost potency both on target and in cells. These compounds compliment the most recent efforts by Reinecke *et al* [35].

These efforts resulted in the next generation of PKN3 inhibitors, with our optimized functional tool compound for PKN3 URS03-06 (**6e**). URS03-06 (**6e**) has good enzyme inhibition against PKN3, potent in cell target engagement (IC_50_=210 nM) and more importantly a narrow kinome profile. This affords URS03-06 (**6e**) the ability to potentially integrate PKN3 biology with appropriate control compounds. In addition the molecular modelling and MD simulations provide new insights into the development of selective chemical probes for PKN3 and beyond and this data set affords a new platform of knowledge on the pyridyl imidazole scaffold previously unavailable to the medicinal chemist’s toolbox.

## Experimental Section

### Modelling methods

Molecular modeling was conducted with Maestro (Schrödinger Releases 2024-2, Maestro, Schrödinger, LLC, New York, NY, 2024) with OPLS4 force field [53].

*Ligand preparation*: small organic molecules were parametrized and minimized using Ligprep module of Schrödinger suite employing OPLS4 force field. Standard settings were obtained to account for potential tautomeric forms around neutral pH. *Protein preparation:* To study complexes between various small organic molecules and the kinase domain of PKN3, we employed our protein model as previously described [36]. In addition, we downloaded the corresponding AlphaFold model (AF-Q6P5Z2-F1-model_v2) for model comparison and validation [54]. The obtained protein structures were prepared with the Protein Preparation Wizard from Schrodinger suite [55]. In this protocol the hydrogens were added, coordination geometry of the H-bond network was optimized, and system was energy minimized by using 0.3 Å heavy atom RMSD convergence. The flexible Induced fit docking was conducted using homology model as protein template for compounds **6a** and **6e**, using analogous approach as previously described [36]. The flexible docking approach was conducted similarly as reported earlier [56]. This flexible docking procedure resulted in a feasible convergence. The visualization of the favorable docking poses was conducted with PyMOL (The PyMOL Molecular Graphics System, Version 3.0.4 Schrödinger, LLC.).

*Molecular dynamics simulations* were carried out using Schrödinger Desmond (Schrödinger Release 2024-2: Desmond Molecular Dynamics System, D. E. Shaw Research, Maestro-Desmond Interoperability Tools, Schrödinger, New York, NY, 2024). In Desmond system builder, the orthorhombic periodic system was created and solvated using TIP3P waters [56]. The systems were further neutralized with a 0.15 M NaCl buffer. At the beginning of the simulation, the system was subjected to the default relaxation protocol of Desmond and heated up to a simulation temperature of 300K. Three unconstrained replicated simulations (randomized seed) up to a length of 1 µs were run using NPT protocol at a temperature of 300 K, pressure of 1.01325 bar, Noe-Hoover thermostat, and timestep of 2 fs. Simulations of 3 replicated trajectories containing 3000 snapshots altogether were combined, aligned and converted to Gromacs xtc-format, after which they were subjected to simulation interactions analysis. Trajectories were analyzed using newer version of Schrodinger Maestro suite (Release 2026-1) and its simulation interactions diagram tools. Visualization of the structures was conducted with PyMOL (The PyMOL Molecular Graphics System, Version 3.1.6.1 Schrödinger, LLC.)

#### Data availability

Full trajectories of the Desmond molecular dynamics simulations are freely available at: https://doi.org/10.5281/zenodo.21934463

### Biology

#### Enzyme Assay and Kinome Profiling

See reference [29] for details and supporting information **Table S1**.

#### NanoBRET method. Cell Transfections and BRET Measurements

*N*-terminal NanoLuc/Kinase fusions were encoded in pFN31K or pFC32K expression vectors (Promega), including flexible Gly-Ser-Ser-Gly linkers between Nluc and each full-length kinase. For cellular BRET target engagement experiments, HEK-293 cells were transfected with NLuc/target fusion constructs using FuGENE HD (Promega) according to the manufacturer’s protocol. Briefly, Nluc/target fusion constructs were diluted into Opti-MEM followed by Transfection Carrier DNA (Promega) at a mass ratio of 1:10 (mass/mass), after which FuGENE HD was added at a ratio of 1:3 (mg DNA: mL FuGENE HD). 1 part (vol) of FuGENE HD complexes thus formed were combined with 20 parts (vol) of HEK-293 cells in DMEM with 10% FBS plated at a density of 2 x 10^5^ per mL into 96-well plates (Corning), followed by incubation in a humidified, 37°C/5% CO_2_ incubator for 20-30 hr. BRET assays were performed in white, 96-well cell culture treated plates (Corning) at a density of 2 x 104 cells/well.

Following transfection, DMEM was exchanged for Opti-MEM. All chemical inhibitors were prepared as concentrated stock solutions in DMSO (Sigma-Aldrich) and diluted in Opti-MEM (unless otherwise noted) to prepare working stocks. Cells were equilibrated for 2 h with energy transfer probes and test compound prior to BRET measurements. Energy transfer probes were prepared at a working concentration of 20X in tracer dilution buffer (12.5 mM HEPES, 31.25% PEG-400, pH 7.5). For target engagement analysis, the energy transfer probes were added to the cells at concentrations optimized for each target. For analysis of PKN3-NL, energy transfer probe K-5 was used at a final concentration of 1000 nM. To measure BRET, NanoBRET NanoGlo Substrate and Extracellular NanoLuc Inhibitor (Promega) were added according to the manufacturer’s recommended protocol, and filtered luminescence was measured on a GloMax Discover luminometer equipped with 450 nM BP filter (donor) and 600 nM LP filter (acceptor), using 0.5 s integration time. Milli-BRET units (mBU) are calculated by multiplying the raw BRET values by 1000. Competitive displacement data were plotted with GraphPad Prism software and data were fit to Equation 1 [log(inhibitor) vs. response -- Variable slope (four parameters)] to determine the IC_50_ value; Y = Bottom + (Top-Bottom)/(1 + 10^((LogIC_50_-X)*HillSlope)). (Equation 1). If the curve was insufficiently described by the raw data, then they were normalized to the controls-BRET in the presence of only energy transfer probe, and BRET in the absence of the energy transfer probe and test compound-then plotted to fit Equation 2 [log(inhibitor) vs. normalized response -- Variable slope] to determine the IC_50_ value; Y=100/(1+10^((LogIC_50_-X)*HillSlope))). (Equation 2). For all BRET data shown, no individual data points were omitted.

### Chemistry

#### General

All reactions were performed using flame-dried round-bottomed flasks or reaction vessels unless otherwise stated. Where appropriate, reactions were carried out under an inert atmosphere of nitrogen with dry solvents, unless otherwise stated. Solvents were obtained directly from the manufacturer and used without further purification. Yields refer to chromatographically and spectroscopically pure isolated yields. Reagents were purchased at the highest commercial-quality and used without further purification, unless otherwise stated. Reactions were monitored by thin-layer chromatography carried out on 0.25 mm E. Merck silica gel plates (60 F254) using ultraviolet light as visualizing agent. NMR spectra were recorded on a Bruker 400 and 600 MHz spectrometers using the residual signal of the deuterated solvent as an internal standard. Coupling constants (J) are given in hertz (Hz). Splitting patterns are described using the following abbreviations or combinations thereof: singlet (s), doublet (d), triplet (t), quartet (q), multiplet (m) and broad singlet (br s). The value of chemical shifts (δ) is given in parts per million (ppm).

#### Route A

##### *Oxidation-Condensation:* 4-(2-(4-Methoxyphenyl)-1*H*-imidazol-5-yl)pyridine (3)

To a stirred solution of 4-acetylpyridine **1** (31.1 g, 257.35 mmol, 1.75 equiv) in DMSO (372.3 mL) was added concentrated aqueous HBr (48% w/w, 8.9 M) (49.57 mL, 441.17 mmol, 3.0 equiv) drop wise at room temperature under air and resulting reaction mixture was stirred at 60 °C for 19 h. After diluting the resulting reaction mixture with MeOH (781.2 mL), was added into the mixture of 4-methoxybenzaldehyde **2** (20.0 g, 147.05 mmol, 1.0 equiv) and NH_4_OAc (226.7 g, 2941.17 mmol, 20.0 equiv) in MeOH (937 mL, 0.16 M in relation to 4-methoxybenzaldehyde) drop wise and was stirred at room temperature for 52h. Reaction mixture was quenched with a mixture of saturated aqueous NaHCO_3_ and saturated aqueous Na_2_S_2_O_3_ solution (1:1, 1000 mL) and was extracted with EtOAc (500 mL × 4). The combined organic extracts were washed with brine (1000 mL), dried over anhydrous Na_2_SO_4_, filtered and concentrated under reduced pressure. The crude product was purified by silica gel (100-200 mesh) column chromatography, using EtOAc: hexanes (gradient elusion 0:10 to 1:9), to afford 4-(2-(4-methoxyphenyl)-1H-imidazol-5-yl)pyridine (**3**) (6.0 g, 9.3%) (6.0 g, 16%) as a yellow solid. ^1^H NMR (400 MHz, DMSO-*d*_*6*_) δ 12.73 (br s, 1H), 8.52 (d, *J* = 5.6 Hz, 2H), 8.01 (s, 1H), 7.95 (d, *J =* 8.4, 2H), 7.78 (d, *J =* 5.6 Hz, 2H), 7.05 (d, *J =* 8.8 Hz, 2H), 3.81 (s, 3H); ^13^C-NMR (101 MHz, DMSO-*d*_*6*_) δ 159.6, 149.8 (2C), 146.8, 141.8, 138.2, 126.6 (2C), 123.0, 118.7 (2C), 116.9, 114.2 (2C), 55.3; MS: [MH]^+^ 251.9

##### *Bromination:* 4-(4-Bromo-2-(4-methoxyphenyl)-1*H*-imidazol-5-yl)pyridine (3I)

To a stirred solution of 4-(2-(4-methoxyphenyl)-1*H*-imidazol-5-yl)pyridine (**3**) (5.0 g, 19.92 mmol, 1.0 equiv) in CH_2_Cl_2_ (75.0 mL, 0.26 M) was added pyridine (21.4 g, 270.91 mmol, 13.6 equiv) followed by bromine (1.23 mL, 3.8 g, 23.90 mmol, 1.2 equiv) drop wise at 0 °C under nitrogen atmosphere and the resulting reaction mixture was stirred at same temperature for 1h. Reaction mixture was dilute with saturated aqueous Na_2_S_2_O_3_ solution (100 mL × 5) and was extracted with 10% MeOH in CH_2_Cl_2_ (100 mL × 4). Combined organic extracts were dried over anhydrous Na_2_SO_4_ and concentrated under reduced pressure. The crude product was purified by silica gel column chromatography, using MeOH: CH_2_Cl_2_ (gradient elusion 0:1 to 1:9) to afford 4-(4-bromo-2-(4-methoxyphenyl)-1*H*-imidazol-5-yl)pyridine (**3I**) (3.4 g, 51%) as a light orange solid. ^1^H NMR (400 MHz, DMSO-*d*_*6*_) δ 12.96 (br s, 1H), 8.67 (d, *J* = 6.1 Hz, 2H), 7.98 (d, *J =* 8.8 Hz, 2H), 7.84 (dd, *J* = 4.7, 1.4 Hz, 1.2 Hz, 2H), 7.08 (d, *J =* 8.8 Hz, 2H), 3.82 (s, 3H); ^13^C-NMR (101 MHz, DMSO-*d*_*6*_) δ 160.2, 150.0 (2C), 147.6, 135.9, 127.2 (2C), 125.1, 121.6, 120.0 (2C), 115.8, 114.3 (2C), 55,3; MS: [MH]^+^ 329.6.

##### *Demethylation:* 4-(4-Bromo-5-(pyridin-4-yl)-1*H*-imidazol-2-yl)phenol (4)

To a stirred solution of 4-(4-bromo-2-(4-methoxyphenyl)-1*H*-imidazol-5-yl)pyridine (**3I**) (3.4 g, 10.33 mmol) in CH_2_Cl_2_ (51.0 mL, 0.2 M) was added 1M BBr_3_ in CH_2_Cl_2_ (41.33 mL, 41.33 mmol, 4.0 equiv) drop wise at 0 °C temperature under nitrogen atmosphere and the resulting reaction mixture was stirred at room temperature for 16h. Reaction mixture was dilute with saturated aqueous NaHCO_3_ solution (100 mL × 1) and was extracted with 20% ^*i*^PrOH in CHCl_3_ (200 mL × 3). The combined organic extracts were dried over anhydrous Na_2_SO_4_ and concentrated under reduced pressure. The crude product was purified by triturating with *n*-pentane (70 mL × 2) and dried under reduced pressure, to afford 4-(4-bromo-5-(pyridin-4-yl)-1H-imidazol-2-yl)phenol (**4**) (2.6 g, 79%) as a light yellow solid. ^1^H NMR (400 MHz, DMSO-*d*_*6*_) δ 12.87 (br s, 1H), 9.84 (br s, 1H), 8.62 (d, *J =* 6.1 Hz, 2H), 7.85-7.83 (m, 4H), 6.85 (dd, *J =* 8.7 Hz, 2H); ^13^C-NMR (101 MHz, DMSO-*d*_*6*_ + one drop of TFA) δ 159.0, 150.7, 145.5, 141.8 (2C), 128.1 (2C), 125.9, 121.0 (2C), 119.6 (2C), 116.0 (2C); MS: [MH]^+^ 315.7.

##### Suzuki-Miyaura cross-coupling

##### Synthesis of 4-(4-(4-Fluorophenyl)-5-(pyridin-4-yl)-1*H*-imidazol-2-yl)phenol (6a)

To a stirred solution of 4-(4-bromo-5-(pyridin-4-yl)-1H-imidazol-2-yl)phenol (**4**) (0.200 g, 0.634 mmol, 1.0 equiv) and (4-fluorophenyl)boronic acid **5a** (0.132 g, 0.952 mmol, 1.5 equiv) in DME: H_2_O (4.5:0.5, 5 mL) was added potassium carbonate (0.306 g, 2.22 mmol, 3.5 equiv) at room temperature under nitrogen and the resulting reaction mixture was degassed (purging with nitrogen) for 15 min followed by the addition of PdCl_2_(PPh_3_)_2_ (0.089 g, 0.126 mmol, 20 mol%) at room temperature. The resulting reaction mixture was irradiated at 150 °C for 30 min in microwave. The reaction mixture was filtered through celite bad and washed with ethyl acetate (25 mL × 4), filtrate was concentrated under reduced pressure, to afford the crude product, which was purified by silica gel column chromatography, using MeOH: CH_2_Cl_2_ (gradient elusion 0:1 to 2:8) to afford ethyl 4-(4-(4-fluorophenyl)-5-(pyridin-4-yl)-1H-imidazol-2-yl)phenol (**6a**) (0.100 g, 47%) as a yellow solid. Melting point: 198-207 °C; ^1^H NMR (400 MHz, DMSO-*d*_*6*_ + one drop of TFA) δ 8.71 (d, *J* = 6.9 Hz, 2H), 7.95 (dd, *J* = 7.8, 2.7 Hz, 4H), 7.72-7.65 (m, 2H), 7.47-7.40 (m, 2H), 6.92 (d, *J* = 8.7 Hz, 2H). (One phenolic proton and one imidazole proton may be exchanged). ^13^C NMR (101 MHz, DMSO-*d*_*6*_) δ 164.3 (^1^*J*_*CF*_ = 247.1 Hz), 160.1, 150.0 (2C), 149.9, 132.1 (^3^*J*_*CF*_ = 8.3 Hz, 2C), 128.7 (2C), 123.1 (2C), 122.2, 116.9 (^2^*J*_*CF*_ = 22.0 Hz, 2C), 116.7 (2C), (4 carbons are missed due to overlapping with other peaks). ^19^F NMR (400 MHz, DMSO-*d*_*6*_ + one drop of TFA) δ −111.4; HRMS m/z [M+H]^+^ calcd for C_20_H_15_FN_3_O: 332.1194, found 332.1206, LC *t*_R_ = 2.63 min, > 98 % purity.

#### Synthesis of 4-(4-(3-fluorophenyl)-5-(pyridin-4-yl)-1*H*-imidazol-2-yl)phenol (6b)

To a stirred solution of 4-(4-bromo-5-(pyridin-4-yl)-1H-imidazol-2-yl)phenol **(4)** (0.200 g, 0.63 mmol, 1.0 equiv) and (3-fluorophenyl)boronic acid **5b** (0.133 g, 0.95 mmol, 1.5 equiv) in DME: H_2_O (4.5:0.5, 5 mL) was added potassium carbonate (0.306 g, 2.22 mmol, 3.5 equiv) at room temperature under nitrogen and the resulting reaction mixture was degassed (purging with nitrogen) for 15 min followed by the addition of PdCl_2_(PPh_3_)_2_ (0.088 g, 0.12 mmol, 20 mol%) at room temperature. The resulting reaction mixture was irradiated at 150 °C for 30 min in microwave. The reaction mixture was filtered through celite bad and washed with ethyl acetate (25 mL × 4), filtrate was concentrated under reduced pressure, to afford the crude product, which was purified by silica gel column chromatography, using MeOH: CH_2_Cl_2_ (gradient elusion 0:1 to 2:8), to afford 4-(4-(3-fluorophenyl)-5-(pyridin-4-yl)-1H-imidazol-2-yl)phenol (**6b**) (0.056 g, 26%) as a pale-yellow solid. Melting point: 178-182 °C; ^1^H NMR (400 MHz, DMSO-*d*_*6*_ + one drop of TFA) δ 8.73 (d, *J* = 6.2 Hz, 2H), 8.01 (d, *J* = 6.3 Hz, 2H), 7.95 (d, *J* = 8.5 Hz, 2H), 7.63-7.55 (m, 1H), 7.53-7.46 (m, 1H), 7.41 (m, 2H), 6.95 (d, *J* = 8.4 Hz, 2H). (One phenolic proton and one imidazole proton may be exchanged); ^13^C-NMR (101 MHz, DMSO-*d*_*6*_ + one drop of TFA) δ 162.5 (^1^*J*_*CF*_ = 245.1 Hz), 159.8, 148.2, 148.1, 142.2 (2C), 134.8, 131.6 (^3^*J*_*CF*_ = 8.5 Hz); 131.4 (^3^*J*_*CF*_ = 8.5 Hz), 129.1, 128.3 (2C), 125.6, 122.6 (2C), 118.3, 116.8 (^2^*J*_*CF*_= 21.0 Hz), 116.3 (^2^*J*_*CF*_ = 22.7 Hz), 116.0 (2C); ^19^F NMR (400 MHz, DMSO-*d*_*6*_ + one drop of TFA) δ −111.7; HRMS m/z [M+H]^+^ calcd for C_20_H_15_FN_3_O: 332.1194, found 332.1210, LC *t*_R_ = 2.66 min, > 98 % purity.

#### Synthesis of 4-(4-(2-fluorophenyl)-5-(pyridin-4-yl)-1*H*-imidazol-2-yl)phenol (6c)

To a stirred solution of 4-(4-bromo-5-(pyridin-4-yl)-1H-imidazol-2-yl)phenol (**4**) (0.250 g, 0.79 mmol, 1.0 equiv) and (2-fluorophenyl)boronic acid **5c** (0.166 g, 1.19 mmol, 1. 5 equiv) in DME: H_2_O (4.5:0.5, 5 mL) was added potassium carbonate (0.383 g, 2.77 mmol, 3.5 equiv) at room temperature under nitrogen and the resulting reaction mixture was degassed (purging with nitrogen) for 15 min followed by the addition of PdCl_2_(PPh_3_)_2_ (0.111 g, 0.15 mmol, 20 mol%) at room temperature. The resulting reaction mixture was irradiated at 150 °C for 35 min in microwave. The reaction mixture was filtered through celite bad and washed with ethyl acetate (25 mL × 4), filtrate was concentrated under reduced pressure. The crude product was purified by (C-18) silica gel column chromatography, using acetonitrile: water (gradient elusion 0:10 to 8:2), to afford 4-(4-(2-fluorophenyl)-5-(pyridin-4-yl)-1H-imidazol-2-yl)phenol (**6c**) (0.060 g, 22%) as a yellow solid. Melting point: 170-180 °C; ^1^H NMR (400 MHz, DMSO-*d*_*6*_ + one drop of TFA) δ 8.73 (d, *J* = 6.9 Hz, 2H), 7.98-7.91 (m, 4H), 7.74-7.63 (m, 2H), 7.49-7.43 (m, 2H), 6.92 (d, *J* = 8.7 Hz, 2H), (One phenolic proton and one imidazole proton may be exchanged); ^13^C NMR (101 MHz, Methanol-*d*_*4*_) δ 161.3 (^1^*J*_*CF*_ = 248.4 Hz), 160.2 (2C), 150.3, 149.7, 143.6, 133.0 (^4^*J*_*CF*_ = 2.0 Hz), 132.2(^3^*J*_*CF*_ = 8.1 Hz), 128.8 (2C), 125.9 (^3^*J*_*CF*_ = 3.7 Hz), 122.1 (2C), 120.9 (^2^*J*_*CF*_ = 15.2 Hz), 117.3 (^2^*J*_*CF*_ = 21.9 Hz), 116.7 (2C); ^19^F NMR (400 MHz, DMSO-*d*_*6*_ + one drop of TFA) δ −112.9; HRMS m/z [M+H]^+^calcd for C_20_H_15_FN_3_O: 332.1194, found 332.1210, LC *t*_R_ = 2.52 min, > 98 % purity.

#### Synthesis of 4-(4-(4-methoxyphenyl)-5-(pyridin-4-yl)-1*H*-imidazol-2-yl)phenol (6d)

To a stirred solution of 4-(4-bromo-5-(pyridin-4-yl)-1H-imidazol-2-yl)phenol (**4**) (0.250 g, 0.79 mmol, 1.0 equiv) and (4-methoxyphenyl)boronic acid **5d** (0.180 g, 1.19 mmol, 1.5 equiv) in DME: H_2_O (4.5:0.5, 5 mL) was added potassium carbonate (0.383 g, 2.77 mmol, 3.5 equiv) at room temperature under nitrogen and the resulting reaction mixture was degassed (purging with nitrogen) for 15 min followed by the addition of PdCl_2_(PPh_3_)_2_ (0.111 g, 0.15 mmol, 20 mol%) at room temperature. The resulting reaction mixture was irradiated at 150 °C for 35min in microwave. The reaction mixture was filtered through celite bad and washed with ethyl acetate (25 mL × 4), filtrate was concentrated under reduced pressure, to afford the crude product, which was purified by silica gel column chromatography, using MeOH: CH_2_Cl_2_ (gradient elusion 0:1 to 2:8), to afford 4-(4-(4-methoxyphenyl)-5-(pyridin-4-yl)-1H-imidazol-2-yl)phenol (**6d**) (0.050 g, 18%) as a pale-yellow solid. Melting point: 293-300 °C; ^1^H NMR (400 MHz, DMSO-*d*_*6*_ + one drop of TFA) δ 8.80 (d, *J* = 6.9 Hz, 2H), 8.02 (d, *J* = 6.9 Hz, 2H), 7.97 (d, *J* = 8.8 Hz, 2H), 7.54 (d, *J* = 8.7 Hz, 2H), 7.11 (d, *J* = 8.8 Hz, 2H), 7.00 (d, *J* = 8.8 Hz, 2H), 3.82 (s, 3H). (One phenolic proton and one imidazole proton may be exchanged); ^13^C NMR (101 MHz, DMSO-*d*_*6*_ + one drop of TFA) δ 160.5, 159.6, 148.0, 147.5, 142.3 (2C), 136.2, 130.8 (2C), 128.5, 128.1 (2C), 121.8 (2C), 121.0, 118.5, 115.9 (2C), 114.8 (2C), 55.4; HRMS m/z [M+H]^+^ calcd for C_21_H_18_N_3_O_2_: 344.1394, found 344.1400, LC *t*_R_ = 2.63 min, > 98 % purity.

#### Synthesis of 4-(4-(3-methoxyphenyl)-5-(pyridin-4-yl)-1*H*-imidazol-2-yl)phenol (6e)

To a stirred solution of 4-(4-bromo-5-(pyridin-4-yl)-1H-imidazol-2-yl)phenol (**4**) (0.300 g, 0.95 mmol, 1.0 equiv) and (3-methoxyphenyl)boronic acid **5e** (0.217 g, 1.42 mmol, 1.5 equiv) in DME: H_2_O (2.7:0.3, 3 mL) was added potassium carbonate (0.394 g, 2.85 mmol, 3.0 equiv) at room temperature under nitrogen and the resulting reaction mixture was degassed (purging with nitrogen) for 15 min followed by the addition of PdCl_2_(PPh_3_)_2_ (0.133 g, 0.19 mmol, 20 mol%) at room temperature. The resulting reaction mixture was irradiated at 150 °C for 35 min in microwave. The Reaction mixture was filtered through celite bad and washed with ethyl acetate (25 mL × 4), filtrate was concentrated under reduced pressure, to afford the crude product, which was purified by silica gel column chromatography, using MeOH: CH_2_Cl_2_ (gradient elusion 0:1 to 2:8), to afford 4-(4-(3-methoxyphenyl)-5-(pyridin-4-yl)-1H-imidazol-2-yl)phenol (**6e**) (0.050 g, 15%) as a pale-yellow solid. Melting point: 240-275 °C; ^1^H NMR (400 MHz, DMSO-*d*_*6*_ + one drop of TFA) δ 8.71 (d, *J* = 6.6 Hz, 2H), 8.00 (d, *J* = 6.6 Hz, 2H), 7.95 (d, *J* = 8.8 Hz, 2H), 7.49 (t, *J* = 7.9 Hz, 1H), 7.21-7.12 (m, 3H), 6.92 (d, *J* = 8.8 Hz, 2H), 3.81 (s, 3H), (One phenolic proton and one imidazole proton may be exchanged); ^13^C NMR (101 MHz, DMSO-*d*_*6*_ + one drop of TFA) δ 160.2, 159.9, 147.8, 147.5, 142.3 (2C), 135.8, 130.8, 129.9, 128.6 (2C), 128.1, 122.6 (2C), 121.5, 117.6, 116.1 (2C), 115.9, 114.8, 55.5; HRMS m/z [M+H]^+^ calcd for C_21_H_18_N_3_O_2_: 344.1394, found 344.1410, LC *t*_R_ = 2.67 min, > 98 % purity.

#### Synthesis of 4-(4-(2-methoxyphenyl)-5-(pyridin-4-yl)-1*H*-imidazol-2-yl)phenol (6f)

To a stirred solution of 4-(4-bromo-5-(pyridin-4-yl)-1H-imidazol-2-yl)phenol **(4)** (0.250 g, 0.79 mmol, 1.0 equiv) and (2-methoxyphenyl)boronic acid **5f** (0.180 g, 1.19 mmol, 1.5 equiv) in DME: H_2_O (4.5:0.5, 5mL) was added potassium carbonate (0.383 g, 2.77 mmol, 3.5 equiv) at room temperature under nitrogen and the resulting reaction mixture was degassed (purging with nitrogen) for 15 min followed by the addition of PdCl_2_(PPh_3_)_2_ (0.111 g, 0.15 mmol, 20 mol%) at room temperature. The resulting reaction mixture was irradiated at 150 °C for 35 min in microwave. The reaction mixture was filtered through celite bad and washed with ethyl acetate (25 mL × 4), filtrate was concentrated under reduced pressure, to afford the crude product, which was purified by silica gel column chromatography, using MeOH: CH_2_Cl_2_ (gradient elusion 0:1 to 2:8), to afford 4-(4-(2-methoxyphenyl)-5-(pyridin-4-yl)-1H-imidazol-2-yl)phenol (**6f**) (0.090 g, 33%) as a pale-yellow solid. Melting point: 266-274 °C; ^1^H NMR (400 MHz, DMSO-*d*_*6*_ + one drop of TFA) δ 8.80 (d, *J* = 6.7 Hz, 2H), 7.98 (d, *J* = 8.7 Hz, 2H), 7.93 (d, *J* = 6.8 Hz, 2H), 7.62 (t, *J* = 8.8 Hz, 1H), 7.53 (dd, *J* = 7.5, 1.5 Hz, 1H), 7.27 (d, *J* = 8.4 Hz, 1H), 7.18 (t, *J*

= 7.56 Hz, 1H), 6.98 (d, *J* = 8.7 Hz, 2H), 3.68 (s, 3H), (One phenolic proton and one imidazole proton may be exchanged); ^13^C NMR (101 MHz, methanol-*d*_*4*_) δ 159.9, 158.8, 149.6 (2C), 149.4, 144.5, 132.6, 131.8, 128.9, 128.6 (2C), 128.3, 122.5, 122.2, 121.9, 121.4, 116.7 (2C), 112.6 (2C), 55.8; HRMS m/z [M+H]^+^ calcd for C_21_H_18_N_3_O_2_: 344.1394, found 344.1426, LC *t*_R_ = 2.48 min, > 98 % purity.

#### Synthesis of 4-(2-(4-hydroxyphenyl)-5-(pyridin-4-yl)-1*H*-imidazol-4-yl)benzonitrile (6g)

To a stirred solution of 4-(4-bromo-5-(pyridin-4-yl)-1H-imidazol-2-yl)phenol (**4**) (0.250 g, 0.79 mmol, 1.0 equiv) and (4-cyanophenyl)boronic acid **5g** (0.175 g, 1.19 mmol, 1.5 equiv) in DME: H_2_O (4.5:0.5, 5 mL) was added potassium carbonate (0.383 g, 2.77 mmol, 3.5 equiv) at room temperature under nitrogen and the resulting reaction mixture was degassed (purging with nitrogen) for 15 min followed by the addition of PdCl_2_(PPh_3_)_2_ (0.111 g, 0.15 mmol, 20 mol%) at room temperature. The resulting reaction mixture was irradiated at 150 °C for 35 min in microwave. The reaction mixture was filtered through celite bad and washed with ethyl acetate (25 mL × 4), filtrate was concentrated under reduced pressure, to afford the crude product, which was purified by silica gel column chromatography, using MeOH: CH_2_Cl_2_ (gradient elusion 0:1 to 2:8), to afford 4-(2-(4-hydroxyphenyl)-5-(pyridin-4-yl)-1H-imidazol-4-yl)benzonitrile (**6g**) (0.060 g, 22%) as a yellow solid. Melting point: 272-280 °C; ^1^H NMR (400 MHz, DMSO-*d*_*6*_) δ 12.82 (br s, 1H), 9.86 (s, 1H), 8.64 (d, *J* = 4.8 Hz, 1H), 8.51 (d, *J* = 5.2 Hz, 1H), 7.96-7.90 (m, 3H), 7.83 (d, *J* = 8.0 Hz, 1H), 7.73 (t, *J* = 7.6 Hz, 2H), 7.50 (d, *J* = 5.3 Hz, 2H), 6.90 (d, *J* = 8.4 Hz, 2H); ^13^C NMR (101 MHz, DMSO-*d*_*6*_ + one drop of TFA) δ 159.8, 148.9, 148.2, 142.2 (2C), 134.8, 134.3, 133.2 (2C), 30.1 (2C), 129.9, 128.3 (2C), 123.0 (2C), 118.7 (2C), 116.0 (2C), 112.2; HRMS m/z [M+H]^+^ calcd for C_21_H_15_N_4_O: 339.1240, found 339.1239, LC *t*_R_ = 2.54 min, > 98 % purity.

#### Synthesis of 3-(2-(4-hydroxyphenyl)-5-(pyridin-4-yl)-1*H*-imidazol-4-yl)benzonitrile (6h)

To a stirred solution of 4-(4-bromo-5-(pyridin-4-yl)-1H-imidazol-2-yl)phenol **(4)** (0.250 g, 0.79 mmol, 1.0 equiv) and (3-cyanophenyl)boronic acid **5h** (0.174 g, 1.19 mmol, 1.5 equiv) in DME: H_2_O (4.5:0.5, 5 mL) was added potassium carbonate (0.328 g, 2.38 mmol, 3.0 equiv) at room temperature under nitrogen and the resulting reaction mixture was degassed (purging with nitrogen) for 15 min followed by the addition of PdCl_2_(PPh_3_)_2_ (0.111 g, 0.15 mmol, 20 mol%) at room temperature. The resulting reaction mixture was irradiated at 150 °C for 35 min in microwave. The reaction mixture was filtered through celite bad and washed with ethyl acetate (25 mL × 4), filtrate was concentrated under reduced pressure, to afford the crude product, which was purified by silica gel column chromatography, using MeOH: CH_2_Cl_2_ (gradient elusion 0:1 to 2:8), to afford 3-(2-(4-hydroxyphenyl)-5-(pyridin-4-yl)-1H-imidazol-4-yl)benzonitrile (**6h**) (0.160 g, 59%) as a yellow solid. Melting point: 276-283 °C; ^1^H NMR (400 MHz, DMSO-*d*_*6*_ + one drop of TFA) δ 8.77 (d, *J* = 6.9 Hz, 2H), 8.12 (t, *J* = 1.4 Hz, 1H), 8.04-7.97 (m, 3H), 7.97-7.94 (m, 2H), 7.91 (dt, *J* = 5.4, 3.8 Hz, 1H), 7.73 (t, *J* = 7.8 Hz, 1H), 6.97 (d, *J* = 8.8 Hz, 2H), (One phenolic proton and one imidazole proton may be exchanged); ^13^C NMR (101 MHz, DMSO-*d*_*6*_ + one drop of TFA) δ 160.0, 148.6, 147.9, 142.3 (2C), 134.3, 134.2, 133.4, 133.0, 130.7, 129.3, 128.4 (2C), 122.9 (2C), 118.4, 118.3, 116.1 (2C), 112.7; HRMS m/z [M+H]^+^ calcd for C_21_H_15_N_4_O: 339.1240, found 339.1249, LC *t*_R_ = 2.50 min, > 98 % purity.

#### Synthesis of 2-(2-(4-Hydroxyphenyl)-5-(pyridin-4-yl)-1*H*-imidazol-4-yl)benzonitrile (6i) and 2-(2-(4-hydroxyphenyl)-5-(pyridin-4-yl)-1*H*-imidazol-4-yl)benzamide (6i’)

To a stirred solution of 4-(4-bromo-5-(pyridin-4-yl)-1H-imidazol-2-yl)phenol (**4**) (0.500 g, 1.58 mmol, 1.0 equiv) and (2-cyanophenyl)boronic acid **5i** (0.349 g, 2.38 mmol, 1.5 equiv) in DME: H_2_O (7.2:0.8, 8 mL) was added potassium carbonate (0.766 g, 5.55 mmol, 3.5 equiv) at room temperature under nitrogen and the resulting reaction mixture was degassed (purging with nitrogen) for 15 min followed by the addition of PdCl_2_(PPh_3_)_2_ (0.222 g, 0.317 mmol, 20 mol%) at room temperature. The resulting reaction mixture was irradiated at 140 °C for 15 min in microwave. The reaction mixture was filtered through celite bad and washed with ethyl acetate (25 mL × 5), filtrate was concentrated under reduced pressure. The crude product was purified by silica gel column chromatography, using MeOH: CH_2_Cl_2_ (gradient elusion 0:1 to 2:8), to afford 2-(2-(4-hydroxyphenyl)-5-(pyridin-4-yl)-1H-imidazol-4-yl)benzonitrile (**6i**) (0.052 g, 5%) as a pale-yellow solid and 2-(2-(4-hydroxyphenyl)-5-(pyridin-4-yl)-1H-imidazol-4-yl)benzamide (**6i’**) as an orange solid which was further purified by (C-18) silica gel column chromatography, using acetonitrile: H_2_O (gradient elusion 0:10 to 4:6), to afford 2-(2-(4-hydroxyphenyl)-5-(pyridin-4-yl)-1H-imidazol-4-yl)benzamide (**6i’**) (0.021 g, 4%) as a pale-yellow solid. (**6i**) Melting point: 264-272 °C; ^1^H NMR (400 MHz, DMSO-*d*_*6*_ + one drop of TFA) δ 8.77 (d, *J* = 6.9 Hz, 2H), 8.12 (s, 1H), 8.01 (m, 3H), 7.98-7.93 (m, 2H), 7.93-7.89 (m, 1H), 7.73 (t, *J* = 7.8 Hz, 1H), 6.99-6.96 (d, *J* = 8.8 Hz, 2H), (One phenolic proton and one imidazole proton may be exchanged); ^13^C NMR (101 MHz, DMSO-*d*_*6*_ + one drop of TFA) δ 159.7, 149.1, 148.7, 142.1 (2C), 134.3 (2C), 133.4 (2C), 133.0, 131.8, 130.9, 128.1 (2C), 121.9 (2C), 119.1, 117.6, 116.1 (2C), 112.5; HRMS m/z [M+H]^+^ calcd for C_21_H_15_N_4_O: 339.1240, found 339.1251, LC *t*_R_ = 2.26 min, > 98 % purity. (**6i’**) Melting point: 273-280 °C; ^1^H NMR (400 MHz, DMSO-*d*_*6*_ + one drop of TFA) δ 8.77 (d, *J* = 6.9 Hz, 2H), 7.98 (d, *J* = 8.7 Hz, 2H), 7.88-7.82 (m, 3H), 7.75-7.66 (m, 2H), 7.59 (dd, *J* = 7.2, 1.5 Hz, 1H), 7.01 (d, *J* = 8.7 Hz, 2H), (One phenolic proton, one imidazole proton and amide protons may be exchanged); ^13^C NMR (101 MHz, DMSO-*d*_*6*_ + one drop of TFA) δ 168.9, 160.2, 147.1, 146.9, 142.1 (2C), 137.5, 135.9, 131.7, 130.9, 130.5, 128.6, 128.4, 128.3 (2C), 127.5, 121.6 (2C), 117.5, 116.1 (2C); HRMS m/z [M+H]^+^ calcd for C_21_H_17_N_4_O_2_: 357.1346, found 357.1348, LC *t*_R_ = 1.86 min, > 98 % purity.

#### Synthesis of 1-(4-(2-(4-Hydroxyphenyl)-5-(pyridin-4-yl)-1*H*-imidazol-4-yl)phenyl)ethan-1-one (6j)

To a stirred solution of 4-(4-bromo-5-(pyridin-4-yl)-1H-imidazol-2-yl)phenol **(4)** (0.250 g, 0.79 mmol, 1.0 equiv) and (4-acetylphenyl)boronic acid **5j** (0.195 g, 1.19 mmol, 1.5 equiv) in DME: H_2_O (4.5:0.5, 5 mL) was added potassium carbonate (0.328 g, 2.38 mmol, 3.0 equiv) at room temperature under nitrogen and the resulting reaction mixture was degassed (purging with nitrogen) for 15 min followed by the addition of PdCl_2_(PPh_3_)_2_ (0.111 g, 0.15 mmol, 20 mol%) at room temperature. The resulting reaction mixture was irradiated at 150 °C for 35 min in microwave. The reaction mixture was filtered through celite bad and washed with ethyl acetate (25 mL × 4), filtrate was concentrated under reduced pressure, to afford the crude product, which was purified by silica gel column chromatography, using MeOH: CH_2_Cl_2_ (gradient elusion 0:1 to 2:8), to afford 1-(4-(2-(4-hydroxyphenyl)-5-(pyridin-4-yl)-1H-imidazol-4-yl)phenyl)ethan-1-one (**6j**) (0.170 g, 60%) as a yellow solid. Melting point: 288-298 °C; ^1^H NMR (400 MHz, DMSO-*d*_*6*_ + one drop of TFA) δ 8.84-8.83 (d, *J*=5.6 Hz, 2H), 8.14-8.12 (d, *J*=8.4 Hz, 2H), 8.05-8.04 (d, *J*=6.4 Hz, 2H), 8.01-7.99 (d, *J*=8.8 Hz, 2H), 7.80-7.78 (d, *J*=8.4 Hz, 2H), 7.00-6.98 (d, *J*=8.0 Hz, 2H), 2.65 (s, 3H), (One phenolic proton and one imidazole proton may be exchanged); ^13^C NMR (101 MHz, DMSO-*d*_*6*_ + one drop of TFA) δ 197.8, 160.2, 148.7, 148.1, 142.5 (2C), 137.7, 135.3, 133.7, 129.8 (2C), 129.4, 129.3 (2C), 128.9 (2C), 123.1 (2C), 118.3, 116.2 (2C), 27.1; HRMS m/z [M+H]^+^ calcd for C_22_H_18_N_3_O_2_: 356.1394, found 339.1398, LC *t*_R_ = 2.50 min, > 98 % purity.

#### Synthesis of 4-(2-(4-hydroxyphenyl)-5-(pyridin-4-yl)-1*H*-imidazol-4-yl)-N-methylbenzamide (6k)

To a stirred solution of 4-(4-bromo-5-(pyridin-4-yl)-1H-imidazol-2-yl)phenol **(4)** (0.200 g, 0.63 mmol, 1.0 equiv) and (4-(methylcarbamoyl)phenyl)boronic acid **5k** (0.170 g, 0.95 mmol) in DME: H_2_O (3.6:0.4, 4 mL) was added potassium carbonate (0.306 g, 2.22 mmol, 3.5 equiv) at room temperature under nitrogen and the resulting reaction mixture was degassed (purging with nitrogen) for 15 min followed by the addition of PdCl_2_(PPh_3_)_2_ (0.088 g, 0.12 mmol, 20 mol%) at room temperature. The resulting reaction mixture was irradiated at 150 °C for 35 min in microwave. The reaction mixture was filtered through celite bad and washed with ethyl acetate (25 mL × 5), filtrate was concentrated under reduced pressure, to afford the crude product, which was purified by silica gel column chromatography, using MeOH: CH_2_Cl_2_ (gradient elusion 0:1 to 2:8), to afford 4-(2-(4-hydroxyphenyl)-5-(pyridin-4-yl)-1H-imidazol-4-yl)-N-methylbenzamide (**6k**) (0.050 g, 21%) as a yellow solid. Melting point: 292-296 °C; ^1^H NMR (400 MHz, DMSO-*d*_*6*_ + one drop of TFA) δ 8.74 (d, *J* = 6.1 Hz, 2H), 8.02-7.94 (m, 6H), 7.70 (d, *J* = 8.2 Hz, 2H), 6.96 (d, *J* = 8.5 Hz, 2H), 2.81 (s, 3H), (One phenolic proton and one imidazole proton may be exchanged); ^13^C NMR (101 MHz, DMSO-*d*_*6*_ + one drop of TFA) δ 166.1, 160.9, 148.1, 146.7, 142.6 (2C), 136.1, 135.1, 130.7, 129.6 (2C), 129.2 (2C), 128.4 (2C), 127.5, 123.5 (2C), 116.5, 116.4 (2C), 26.6; HRMS m/z [M+H]^+^ calcd for C_22_H_19_N_4_O_2_: 371.1503, found 371.1502, LC *t*_R_ = 2.04 min, > 98 % purity.

#### Synthesis of N-(4-(2-(4-hydroxyphenyl)-5-(pyridin-4-yl)-1*H*-imidazol-4-yl)phenyl)acetamide (6l)

To a stirred solution of 4-(4-bromo-5-(pyridin-4-yl)-1H-imidazol-2-yl)phenol (**4**) (0.220 g, 0.69 mmol, 1.0 equiv) and (4-acetamidophenyl)boronic acid **5l** (0.187 g, 1.04 mmol, 1.5 equiv) in DME: H_2_O (4.5:0.5, 5 mL) was added potassium carbonate (0.289 g, 2.09 mmol, 3.0 equiv) at room temperature under nitrogen and the resulting reaction mixture was degassed (purging with nitrogen) for 15 min followed by the addition of PdCl_2_(PPh_3_)_2_ (0.098 g, 0.13 mmol, 20 mol%) at room temperature. The resulting reaction mixture was irradiated at 150 °C for 35 min in microwave. The reaction mixture was filtered through celite bad and washed with ethyl acetate (25 mL × 4), filtrate was concentrated under reduced pressure, to afford the crude product, which was purified by silica gel column chromatography, using MeOH: CH_2_Cl_2_(gradient elusion 0:1 to 2:8), to afford N-(4-(2-(4-hydroxyphenyl)-5-(pyridin-4-yl)-1H-imidazol-4-yl)phenyl)acetamide (**6l**) (0.135 g, 52%) as a yellow solid. Melting point: 284-294 °C; ^1^H NMR (400 MHz, DMSO-*d*_*6*_ + one drop of TFA) δ 8.72 (d, *J* = 6.8 Hz, 2H), 8.00 (d, *J* = 6.9 Hz, 2H), 7.95 (d, *J* = 8.7 Hz, 2H), 7.75 (d, *J* = 8.6 Hz, 2H), 7.53 (d, *J* = 8.6 Hz, 2H), 6.96 (d, *J* = 8.7 Hz, 2H), 2.08 (s, 3H), (One phenolic proton and one imidazole proton may be exchanged); ^13^C NMR (101 MHz, DMSO-*d*_*6*_ + one drop of TFA) δ 169.1, 160.5, 147.5, 146.9, 142.4 (2C), 141.3, 135.9, 130.1 (2C), 128.9 (2C), 126.9, 122.8 (2C), 122.2, 119.5 (2C), 116.2, 116.7 (2C), 24.2; HRMS m/z [M+H]^+^ calcd for C_22_H_19_N_4_O_2_: 371.1503, found 371.1502, LC *t*_R_ = 2.13 min, > 98 % purity.

#### Synthesis of 4-(4-(4-(methylsulfonyl)phenyl)-5-(pyridin-4-yl)-1*H*-imidazol-2-yl)phenol (6m)

To a stirred solution of 4-(4-bromo-5-(pyridin-4-yl)-1H-imidazol-2-yl)phenol (**4**) (0.220 g, 0.69 mmol, 1.0 equiv) and (4-

(methylsulfonyl)phenyl)boronic acid **5m** (0.209 g, 1.04 mmol, 1.5 equiv) in DME: H_2_O (4.5:0.5, 5 mL) was added potassium carbonate (0.289 g, 2.09 mmol, 3.0 equiv) at room temperature under nitrogen and the resulting reaction mixture was degassed (purging with nitrogen) for 15 min followed by the addition of PdCl_2_(PPh_3_)_2_ (0.098 g, 0.13 mmol, 20 mol%) at room temperature. The resulting reaction mixture was irradiated at 150 °C for 35 min in microwave. The reaction mixture was filtered through celite bad and washed with ethyl acetate (25 mL × 4), filtrate was concentrated under reduced pressure, to afford the crude product, which was purified by silica gel column chromatography, using MeOH: CH_2_Cl_2_ (gradient elusion 0:1 to 2:8), to afford 4-(4-(4-(methylsulfonyl)phenyl)-5-(pyridin-4-yl)-1H-imidazol-2-yl)phenol **(6m)** (0.137 g, 50%) as a pale-yellow solid. Melting point: 286-290 °C; ^1^H NMR (400 MHz, DMSO-*d*_*6*_ + one drop of TFA) δ 8.74 (d, *J* = 6.2 Hz, 2H), 8.05 (m, 4H), 7.95 (d, *J* = 8.5 Hz, 2H), 7.87 (d, *J* = 8.2 Hz, 2H), 6.96 (d, *J* = 8.5 Hz, 2H), 3.26 (s, 3H), (One phenolic proton and one imidazole proton may be exchanged); ^13^C NMR (101 MHz, DMSO-*d*_*6*_ + one drop of TFA) δ 160.3, 148.7, 147.5, 142.4 (2C), 141.8, 134.6, 134.1, 130.3 (2C), 129.1, 128.7 (2C), 128.1 (2C), 123.4 (2C), 117.8, 116.2 (2C), 43.6; HRMS m/z [M+H]^+^ calcd for C_21_H_18_N_3_O_3_S: 392.1063, found 392.1075, LC *t*_R_ = 2.25 min, > 98 % purity.

#### Phenol alkylation

##### Synthesis of 1-(4-(4-(4-Fluorophenyl)-5-(pyridin-4-yl)-1*H*-imidazol-2-yl)phenoxy)-2-methylpropan-2-ol (6n)

To a stirred solution of 4-(4-(4-fluorophenyl)-5-(pyridin-4-yl)-1H-imidazol-2-yl)phenol **(6a)** (0.200 g, 0.60 mmol) and 2,2-dimethyloxirane (0.2 mL, 2.25 mmol, 3.75 equiv) in DMF (2 mL) was added NaHCO_3_ (0.177 g, 2.11 mmol, 3.5 equiv) at room temperature under nitrogen and the resulting reaction mixture was irradiated at 150 °C for 90 min in microwave. The reaction mixture was diluted with H_2_O (80 mL) and was extracted with EtOAc (100 mL × 2). The combined organic extracts were dried over anhydrous Na_2_SO_4_ and concentrated under reduced pressure. The crude product was purified by prep HPLC, using 5 mmol (NH_4_)HCO_3_ in H_2_O: MeCN, to afford 1-(4-(4-(4-fluorophenyl)-5-(pyridin-4-yl)-1H-imidazol-2-yl)phenoxy)-2-methylpropan-2-ol (**6n**) (0.040 g, 16%) as a pale yellow solid. Melting point: 150-158 °C; ^1^H NMR (400 MHz, DMSO-*d*_*6*_ + one drop of TFA) δ 8.65 (d, *J* = 6.9 Hz, 2H), 8.04-7.98 (m, 4H), 7.68-7.64 (m, 2H), 7.40 (t, *J* = 8.8 Hz, 2H), 7.11 (d, *J* = 8.8 Hz, 2H), 3.78 (s, 2H), 1.21 (s, 6H), (One imidazole proton and one aliphatic hydroxy proton may be exchanged); ^13^C NMR (101 MHz, DMSO-*d*_*6*_ + one drop of TFA) δ 162.9 (^1^*J*_*CF*_ = 247.4 Hz), 160.3, 149.1, 147.7, 141.9 (2C), 135.8, 131.7 (^3^*J*_*CF*_ = 8.6 Hz, 2C), 130.0, 127.6 (2C), 126.1 (^4^*J*_*CF*_ = 2.6 Hz), 121.9 (2C), 120.9, 116.4 (^2^*J*_*CF*_ = 21.8 Hz, 2C), 115.1 (2C), 76.3, 68.7, 26.6 (2C); ^19^F NMR (400 MHz, DMSO-*d*_*6*_ + one drop of TFA) δ −111.3; HRMS m/z [M+H]^+^ calcd for C_24_H_23_FN_3_O_2_: 404.1769, found 404.1790, LC *t*_R_ = 3.32 min, > 98 % purity.

#### Synthesis of 4-(4-(4-fluorophenyl)-2-(4-(2-methoxyethoxy)phenyl)-1*H*-imidazol-5-yl)pyridine (6o)

To a stirred solution of 4-(4-(4-fluorophenyl)-5-(pyridin-4-yl)-1H-imidazol-2-yl)phenol (**6a**) (0.200 g, 0.60 mmol) and 1-bromo-2-methoxyethane (0.417 g, 3.00 mmol, 5 equiv) in DMF (4 mL) was added K_2_CO_3_ (0.415 g, 3.00 mmol, 5 equiv) at room temperature under nitrogen and the resulting reaction mixture was degassed (purging with nitrogen) for 15 min and the resulting reaction mixture was heated at 60 °C for 3h. Reaction mixture was dilute with H_2_O (50 mL) and was extracted with EtOAc (100 mL × 2). The combined organic extracts were washed with brine (100 mL) were dried over anhydrous Na_2_SO_4_ and concentrated under reduced pressure. The crude product was purified by silica gel column chromatography, using MeOH: CH_2_Cl_2_ (gradient elusion 0:1 to 2:8), to afford 4-(4-(4-fluorophenyl)-2-(4-(2-methoxyethoxy)phenyl)-1H-imidazol-5-yl)pyridine **(6o)** (0.100 g, 42%) as a yellow solid. Melting point: 190-200 °C; ^1^H NMR (400 MHz, DMSO-*d*_*6*_ + one drop of TFA) δ 8.71 (d, *J* = 7.0 Hz, 2H), 8.07-8.02 (m, 2H), 8.00 (d, *J* = 7.0 Hz, 2H), 7.72-7.64 (m, 2H), 7.46-7.38 (m, 2H), 7.16-7.10 (m, 2H), 4.19-4.16 (m, 2H), 3.69-3.67 (m, 2H), 3.31 (s, 3H). (One imidazole proton may be exchanged). ^13^C NMR (100 MHz, DMSO-*d*_*6*_ + one drop of TFA) δ 162.8 (^1^*J*_*CF*_ *=* 247.3 Hz), 159.8, 149.2, 147.6, 141.9 (2C), 135.8, 131.7 (^3^*J*_*CF*_ *=* 8.7 Hz, 2C), 130.2, 127.5 (2C), 126.2, 121.8 (2C), 121.3, 116.4 (^2^*J*_*CF*_ *=* 21.8 Hz, 2C), 114.9 (2C), 70.4, 67.2, 58.3; ^19^F NMR (400 MHz, DMSO-*d*_*6*_ + one drop of TFA) δ −111.2; HRMS m/z [M+H]^+^ calcd for C_23_H_21_FN_3_O_2_: 390.1612, found 390.1618, LC *t*_R_ = 3.35 min, > 98 % purity.

#### Route B

##### 1-(4-Fluorophenyl)-2-(pyridin-4-yl) ethan-1-one (9)

*Preparation of LDA solution:* A solution of LDA was prepared by adding a 2.5 M solution of n-butyl lithium (128.4 mL, 321.0 mmol) to a stirred solution of N,N’-diisopropylamine (32.42 g, 321.0 mmol) in dry THF (200 mL) at −78 °C under nitrogen atmosphere and the mixture was stirred at the same temperature for 30 min. The LDA solution (1.5 equiv) was then added drop wise to a solution of 4-methylpyridine **8** (20.0 g, 214 mmol) in THF (100 mL) at −78 °C. After stirring of 15 min, a solution of 4-fluoro-N-methoxy-N-methylbenzamide **7** (39.20 g, 214 ml) was added in THF (100 mL) into the reaction mixture at same temperature and resulting mixture was stirred at 0 °C for 30 min. The reaction was quenched with H_2_O (200 mL) at 0 °C and concentrated HCl (pH~6) and extracted with Et_2_O (3 x 200 mL). The combined organic extracts were dried over anhydrous Na_2_SO_4_ and concentrated under reduced pressure. The obtained crude was purified by silica gel (60-120 mesh) column chromatography, using EtOAc:Hexanes (gradient elusion 0:1 to 5:5), to afford methyl 1-(4-fluorophenyl)-2-(pyridin-4-yl)ethan-1-one (**9**) (35.0 g, 76%) as a white solid. ^1^H NMR (400 MHz, DMSO) δ 8.52-8.50 (dd, *J*=4.4 Hz, 1,6 Hz, 2H), 8.16-8.12 (m, 2H), 7.41-7.38 (m, 2H), 7.30-7.28 (m, 2H), 4.49 (s, 2H); MS: [MH]^+^ 215.9

##### (E)-1-(4-fluorophenyl)-2-(hydroxyimino)-2-(pyridin-4-yl)ethan-1-one (10)

To a stirred solution of 1-(4-fluorophenyl)-2-(pyridin-4-yl)ethan-1-one **9** (33.0 g, 153.0 mmol) in AcOH (150 mL) was added of aqueous solution of NaNO_2_ (0.1 M) (2.3 L, 230 mmol, 1.5 equiv) dropwise into the reaction mixture and the resulting reaction mixture was stirred at 0 °C for 15 min. After completion of reaction, the resulting precipitate was filtered and dried under reduced pressure. The obtained crude was purified by silica gel (60-120 mesh) column chromatography, using EtOAc:Hexanes (gradient elusion 0:1 to 5:5) to afford (E)-1-(4-fluorophenyl)-2-(hydroxyimino)-2-(pyridin-4-yl)ethan-1-one (**10**) (40.5 g, quantitative) as a white solid. ^1^H NMR (400 MHz, DMSO) δ 12.47 (s, 1H), 8.63-8.62 (dd, *J*=4.4 Hz, 1.6 Hz, 2H)), 7.96-7.92 (m, 2H), 7.74-7.41 (m, 4H); MS: [MH]^+^ 244.9

##### 2-(4-Bromophenyl)-4-(4-fluorophenyl)-5-(pyridin-4-yl)-1*H*-imidazol-1-ol (10I)

To a stirred solution of (E)-1-(4-fluorophenyl)-2-(hydroxyimino)-2-(pyridin-4-yl)ethan-1-one (**10**) (20.0 g, 81.96 mmol) in acetic acid (200 mL) were added 4-bromobenzaldehyde **11** (19.71 g, 106.55 mmol, 1.3 equiv) and NH_4_OAc (41.02 g, 532.78 mmol, 6.5 equiv) portion wise at room temperature. The resulting reaction mixture was stirred at 110 °C for 16h. Reaction mixture was cooled to room temperature, basified (pH ~ 6-7) with aqueous NH_4_OH solution and resulting precipitate was collected by filtration. The obtained crude was purified by silica gel (60-120 mesh) column chromatography, using MeOH: CH_2_Cl_2_ (gradient elusion 0:1 to 2:8) to afford, 2-(4-bromophenyl)-4-(4-fluorophenyl)-5-(pyridin-4-yl)-1H-imidazol-1-ol (**10I**) (23.0 g, 68%) as a brown solid. ^1^H NMR (400 MHz, DMSO) δ 12.39 (bs, 1H), 8.64 (d, *J* = 5.9 Hz, 2H), 8.13-8.04 (m, 2H), 7.75-7.68 (m, 2H), 7.54-7.43 (m, 4H), 7.22-7.12 (m, 2H); ^13^C NMR (100 MHz, DMSO-*d*_*6*_) δ 161.5 (^1^*J*_*CF*_ = 244.2 Hz), 149.9 (2C), 140.4, 136.3, 133.8, 131.6 (2C), 130.4, 129.0 (^3^*J*_*CF*_ = 8.2 Hz, 2C), 128.9 (2C), 127.9, 125.0, 124.0 (2C), 122.2, 115.4 (^2^*J*_*CF*_ = 21.4 Hz, 2C); MS: [MH]^+^ 410.1

##### 4-(2-(4-Bromophenyl)-4-(4-fluorophenyl)-1*H*-imidazol-5-yl)pyridine (12)

To a stirred solution of 2-(4-bromophenyl)-4-(4-fluorophenyl)-5-(pyridin-4-yl)-1H-imidazol-1-ol (**11I**) (10.0 g, 24.44 mmol) in DMF (100 mL) was added P(OEt)_3_ (4.05 g, 24.44 mmol, 1.0 equiv) at room temperature under nitrogen atmosphere and the resulting reaction mixture was stirred at 110 °C for 16h. Reaction mixture was diluted with H_2_O (300 mL) and extracted with EtOAc (300 mL × 4). The combined organic extracts were washed with brine (100 mL), dried over anhydrous Na_2_SO_4_ and concentrated under reduced pressure, to afford 4-(2-(4-bromophenyl)-4-(4-fluorophenyl)-1H-imidazol-5-yl)pyridine (**12**) (9.0 g, 93%) as a yellow solid. ^1^H NMR (400 MHz, DMSO) δ 13.02 (br s, 1H), 8.52 (d, *J*=5.5 Hz, 2H), 8.08-7.97 (m, 2H), 7.78-7.69 (m, 2H), 7.63-7.53 (m, 2H), 7.50-7.43 (m, 2H), 7.30 (t, *J*=8.8 Hz, 2H); MS: [MH]^+^ 393.7

#### Buchwald-Hartwig cross-coupling

##### 1-(4-(4-(4-fluorophenyl)-5-(pyridin-4-yl)-1*H*-imidazol-2-yl)phenyl)-4-methylpiperazine (14a)

To a stirred solution of 4-(2-(4-bromophenyl)-4-(4-fluorophenyl)-1H-imidazol-5-yl)pyridine **(12)** (0.500 g, 1.27 mmol) in t-BuOH: 1,4-dioxane (1:1.5 mL) were added 1-methylpiperazine **13a** (0.890 g, 8.90 mmol, 7.0 equiv) and NaO^t^Bu (0.427 g, 4.45 mmol, 3.5 equiv) at room temperature under nitrogen and the resulting reaction mixture was degassed (purging with nitrogen) for 15 min followed by the addition of DavePhos (0.099 g, 0.254 mmol, 20 mol%) and Pd_2_(dba)_3_ (0.116 g, 0.127 mmol, 10 mol%) at room temperature. The resulting reaction mixture was irradiated at 120 °C for 1h in microwave. The reaction mixture was diluted with H_2_O (50 mL) and was extracted with EtOAc (50 mL × 3). The combined organic extracts were dried over anhydrous Na_2_SO_4_ and concentrated under reduced pressure.The crude product was purified by (C-18) silica gel column chromatography, using MeCN:H_2_O (gradient elusion 0:10 to 8:2), to afford 1-(4-(4-(4-fluorophenyl)-5-(pyridin-4-yl)-1H-imidazol-2-yl)phenyl)-4-methylpiperazine (**14a**) (0.070 g, 13%) as a yellow solid. Melting point: 271-280 °C; ^1^H NMR (400 MHz, DMSO-*d*_*6*_, High temperature) δ 12.43 (s, 1H), 8.46 (s, 2H), 7.93 (d, *J* = 8.9 Hz, 2H), 7.60 – 7.53 (m, 2H), 7.47 (d, *J* = 5.2 Hz, 2H), 7.27 (s, 2H), 7.01 (d, *J* = 9.0 Hz, 2H), 3.27 – 3.22 (m, 4H), 2.48 – 2.47 (m, 4H), 2.25 (s, 3H). ^13^C NMR (400 methanol-*d*_*4*_) δ 153.0, 150.1 (2C), 132.11 (^3^*J*_*CF*_ = 8.2 Hz, 2C), 132.1, 128.2 (2C), 123.1 (2C), 121.9, 117.0, 116.9 (^2^*J*_*CF*_ = 21.4 Hz, 2C), 55.7, 45.7; ^19^F NMR (400 MHz, DMSO-*d*_*6*_ + one drop of TFA) δ 110.8 Hz; HRMS m/z [M+H]^+^ calcd for C_25_H_25_FN_5_: 414.2089, found 414.2092, LC *t*_R_ = 2.05 min, > 98 % purity.

##### 4-(4-(4-(4-Fluorophenyl)-5-(pyridin-4-yl)-1*H*-imidazol-2-yl)phenyl)morpholine (14b)

To a stirred solution of 4-(2-(4-bromophenyl)-4-(4-fluorophenyl)-1H-imidazol-5-yl)pyridine (**12**) (0.500 g, 1.27 mmol) and morpholine **13b** (0.774 g, 8.90 mmol, 7.0 equiv) in t-BuOH-1,4-dioxane (1:1.5 mL) was added NaO^t^Bu (0.427 g, 4.45 mmol, 3.5 equiv) at room temperature under nitrogen and the resulting reaction mixture was degassed (purging with nitrogen) for 15 min followed by the addition of DavePhos (0.099 g, 0.254 mmol, 20 mol%) and Pd_2_(dba)_3_ (0.116 g, 0.127 mmol, 10 mol%) at room temperature. The resulting reaction mixture was irradiated at 120 °C for 1h in microwave. The reaction mixture was diluted with H_2_O (50 mL) and was extracted with 20% ^*i*^PrOH:CHCl_3_ (50 mL × 3). The combined organic extracts were dried over anhydrous Na_2_SO_4_ and concentrated under reduced pressure. The crude product was purified by prep HPLC, using 5 mmol (NH_4_)HCO_3_ and 0.05% NH_4_OH in H_2_O:MeCN, to afford 4-(4-(4-(4-fluorophenyl)-5-(pyridin-4-yl)-1H-imidazol-2-yl)phenyl)morpholine (**14b**) (0.035 g, 7%) as a white solid. Melting point: above 300 °C; ^1^H NMR (400 MHz, DMSO-*d*_*6*_ + one drop of TFA) δ 8.45 (d, *J* = 5.7 Hz, 2H), 7.95 – 7.87 (m, 2H), 7.55 – 7.47 (m, 2H), 7.43 (d, *J* = 5.5 Hz, 2H), 7.25 (t, *J* = 8.6 Hz, 2H), 7.02 (d, *J* = 8.8 Hz, 2H), 3.75 – 3.66 (m, 4H), 3.17 (s, 4H), (One imidazole proton may be exchanged); ^13^C NMR (100 MHz, DMSO-*d*_*6*_ + one drop of TFA) δ 163.0 (^1^*J*_*CF*_ = 247 Hz), 152.2, 147.7, 142.4 (2C), 135.1, 131.8 (^3^*J*_*CF*_ = 8.5 Hz, 2C), 128.6, 127.5 (2C), 125.4, 122.2 (2C), 116.9, 116.5 (^2^*J*_*CF*_ = 21.9 Hz, 2C), 114.4 (2C), 66.0 (2C), 47.4 (2C).); ^19^F NMR (400 MHz, DMSO-*d*_*6*_ + one drop of TFA) δ −110.1 Hz; HRMS m/z [M+H]^+^ calcd for C_24_H_22_FN_4_O: 401.1772, found 401.1769, LC *t*_R_ = 3.22 min, > 98 % purity.

#### 1-(4-(4-(4-Fluorophenyl)-5-(pyridin-4-yl)-1*H*-imidazol-2-yl)phenyl)piperazine (14c)

To a stirred solution of 4-(2-(4-bromophenyl)-4-(4-fluorophenyl)-1H-imidazol-5-yl)pyridine (**12**) (0.500 g, 1.27 mmol) and piperazine **13c** (0.765 g, 8.90 mmol, 7.0 equiv) in dry 1,4-dioxane (5 mL) was added NaO^t^Bu (0.427 g, 4.45 mmol, 3.5 equiv) at room temperature under nitrogen and the resulting reaction mixture was degassed (purging with nitrogen) for 15 min followed by the addition of DavePhos (0.099 g, 0.254 mmol, 20 mol%) and Pd_2_(dba)3 (0.058 g, 0.063 mmol, 5 mol%) at room temperature. The resulting reaction mixture was stirred at 120 °C for 1h in microwave. The reaction mixture was diluted with H2O (100 mL) and was extracted with EtOAc (200 mL × 2). The combined organic extracts were dried over anhydrous Na_2_SO_4_ and concentrated under reduced pressure. The crude product was purified by Prep HPLC, using 5 mmol (NH_4_)HCO_3_ in H_2_O:MeCN, to afford 1-(4-(4-(4-fluorophenyl)-5-(pyridin-4-yl)-1H-imidazol-2-yl)phenyl)piperazine (**14c**) (0.080 g, 15%) as a white solid. Melting point: 257-270 °C; ^1^H NMR (400 MHz, DMSO-*d*_*6*_ + one drop of TFA) δ δ 8.86 (s, 2H), 8.78 (d, *J* = 6.6 Hz, 2H), 8.04 (d, *J* = 8.9 Hz, 2H), 7.99 (d, *J* = 6.8 Hz, 1H), 7.69 (dd, *J* = 8.6, 5.4 Hz, 2H), 7.43 (t, *J* = 8.8 Hz, 2H), 7.19 (d, *J* = 8.9 Hz, 2H), 3.53 (d, *J* = 5.2 Hz, 4H), 3.26 (s, 4H); ^13^C NMR (100 MHz, DMSO-*d*_*6*_ + one drop of TFA) δ δ 163.3 (^1^*J*_*CF*_ = 248.2 Hz), 151.5, 147.6, 147.3, 142.4 (2C), 135.2, 132.0 (^3^*J*_*CF*_ = 8.6 Hz, 2C), 128.1, 127.9 (2C), 125.1, 122.7 (2C), 117.1, 116.62 (d, ^2^*J*_*CF*_ = 21.9 Hz, 2C), 115.3 (2C), 44.6 (2C), 42.8 (2C); ^19^F NMR (400 MHz, DMSO-*d*_*6*_ + one drop of TFA) δ −111.1 Hz; HRMS m/z [M+H]^+^ calcd for C_24_H_23_FN_5_: 400.1932, found 400.1924, LC *t*_R_ = 2.02 min, > 98 % purity.

#### N^1^-(4-(4-(4-fluorophenyl)-5-(pyridin-4-yl)-1*H*-imidazol-2-yl)phenyl)-N^1^,N^2^,N^2^-trimethylethane-1,2-diamine (14d)

To a stirred solution of 4-(2-(4-bromophenyl)-4-(4-fluorophenyl)-1H-imidazol-5-yl)pyridine (**12**) (0.300 g, 0.763 mmol) and N^1^,N^1^,N^2^-trimethylethane-1,2-diamine **13d** (0.077 g, 0.763 mmol, 1.0 equiv) in DMA (3 mL) was added NaO^t^Bu (0.256 g, 2.67 mmol, 3.5 equiv) at room temperature under nitrogen and the resulting reaction mixture was degassed (purging with nitrogen) for 15 min followed by the addition of RuPhos (0.071 g, 0.152 mmol, 20 mol%) and Pd_2_(dba)_3_ (0.034 g, 0.038 mmol, 5 mol%) at room temperature. The resulting reaction mixture was irradiated at 150 °C for 45 min in microwave. Reaction mixture was diluted with H_2_O (50 mL) and was extracted with EtOAc (50 mL × 2). The combined organic extracts were washed with brine (50 mL), dried over anhydrous Na_2_SO_4_ and concentrated under reduced pressure. The crude product was purified by (C-18) silica gel column chromatography, using MeCN:H_2_O (gradient elusion 0:10 to 6:5), to afford N^1^-(4-(4-(4-fluorophenyl)-5-(pyridin-4-yl)-1H-imidazol-2-yl)phenyl)-N^1^,N^2^,N^2^-trimethylethane-1,2-diamine (**14d**) (0.056 g, 17%) as a yellow solid. Melting point: 110-115 °C; ^1^H NMR (400 MHz, DMSO-*d*_*6*_ + one drop of TFA) δ 8.75 (d, *J* = 6.9 Hz, 2H), 7.98 (dd, *J* = 8.0, 3.8 Hz, 4H), 7.66 (dd, *J* = 8.7, 5.4 Hz, 2H), 7.40 (t, *J* = 8.8 Hz, 2H), 6.95 (d, *J* = 9.1 Hz, 2H), 3.84 – 3.70 (m, 2H), 3.32 – 3.19 (m, 2H), 3.01 (s, 3H), 2.84 (s, 6H), (One imidazole proton may be exchanged); ^13^C NMR (100 MHz, DMSO-*d*_*6*_ + one drop of TFA) δ 162.8 (^1^*J*_*CF*_ = 247.2 Hz), 149.7, 147.9, 143.0, 134.9, 131.7 (^3^*J*_*CF*_=8.5 Hz, 2C), 129.1, 127.5 (2C), 126.0, 121.7 (2C), 116.3 (^2^*J*_*CF*_ = 21.9 Hz, 2C), 115.8,, 112.1 (2C), 52.6, 46.4, 42.7(2C), 38.0.; ^19^F NMR (400 MHz, DMSO-*d*_*6*_ + one drop of TFA) δ −110.5 Hz; HRMS m/z [M+H]^+^ calcd for C_25_H_27_FN_5_: 416.2245, found 416.2254, LC *t*_R_ = 2.20 min, > 98 % purity.

#### (R)-1-(4-(4-(4-fluorophenyl)-5-(pyridin-4-yl)-1*H*-imidazol-2-yl)phenyl)-2,4-dimethylpiperazine (14e)

To a stirred solution of 4-(2-(4-bromophenyl)-4-(4-fluorophenyl)-1H-imidazol-5-yl)pyridine (**12**) (0.400 g, 1.01 mmol) and (*R*)-1,3-dimethylpiperazine **13e** (203 μL, 1.52 mmol, 1.5 equiv) in DMA (4 mL) was added NaO^t^Bu (0.341 g, 3.56 mmol, 3.5 equiv) at room temperature under nitrogen and the resulting reaction mixture was degassed (purging with nitrogen) for 15 min followed by the addition of RuPhos (0.094 g, 0.203 mmol, 20 mol%) and Pd_2_(dba)_3_ (0.046 g, 0.050 mmol, 5 mol%) at room temperature. The resulting reaction mixture was irradiated at 150 °C for 1h in microwave. The reaction mixture was diluted with H_2_O (100 mL) and was extracted with EtOAc (100 mL × 3). The combined organic extracts were washed with brine, dried over anhydrous Na_2_SO_4_ and concentrated under reduced pressure. The crude product was purified by silica gel column chromatography, using MeOH:CH_2_Cl_2_ (gradient elusion 0:1 to 2:8), followed by (C-18) silica gel column chromatography, using MeCN:H_2_O (gradient elusion 0:10 to 2:8), to afford (R)-1-(4-(4-(4-fluorophenyl)-5-(pyridin-4-yl)-1H-imidazol-2-yl)phenyl)-2,4-dimethylpiperazine (**14e**) (0.050 g, 14%) as a pale-yellow solid. Melting point: 264-268 °C; ^1^H NMR (400 MHz, DMSO-*d*_*6*_ + one drop of TFA) δ 8.79 (d, *J* = 6.6 Hz, 2H), 8.01 (dd, *J* = 11.2, 7.9 Hz, 4H), 7.60 (dd, *J* = 8.5, 5.2 Hz, 2H), 7.27 (t, *J* = 8.7 Hz, 2H), 7.11 (d, *J* = 8.9 Hz, 2H), 4.51 (br s, 1H), 3.88 (d, *J* = 12.8 Hz, 1H), 3.59-3.40 (m, 2H), 3.35-3.20 (m, 2H), 3.13-3.02 (m, 1H), 2.83 (s, 3H), 1.18 (d, *J* = 6.7 Hz, 3H), (One imidazole proton may be exchanged); ^13^C NMR (100 MHz, DMSO-*d*_*6*_ + one drop of TFA) δ 162.7 (^1^*J*_*CF*_ = 247 Hz), 149.2, 147.72, 142.6, 131.6 (^3^*J*_*CF*_ = 8.6 Hz, 2C), 127.2 (2C), 126.3, 121.7 (2C), 116.3 (^2^*J*_*CF*_ = 21.9 Hz, 2C),114.9, 56.6, 52.6, 47.6, 43.0, 38.3, 11.6.; ^19^F NMR (400 MHz, DMSO-*d*_*6*_ + one drop of TFA) δ − 111.3 Hz; HRMS m/z [M+H]^+^ calcd for C_26_H_27_FN_5_: 428.2245, found 428.2247, LC *t*_R_ = 2.19 min, > 98 % purity.

#### (S)-1-(4-(4-(4-fluorophenyl)-5-(pyridin-4-yl)-1*H*-imidazol-2-yl)phenyl)-2,4-dimethylpiperazine (14f)

To a stirred solution of 4-(2-(4-bromophenyl)-4-(4-fluorophenyl)-1H-imidazol-5-yl)pyridine (**12**) (0.700 g, 1.78 mmol) and (S)-1,3-dimethylpiperazine **13f** (0.347 μL,, 2.67 mmol, 1.5 equiv) in DMA (7 mL) was added NaO^t^Bu (0.598 g, 6.23 mmol, 3.5 equiv) at room temperature under nitrogen and the resulting reaction mixture was degassed (purging with nitrogen) for 20 min followed by the addition of RuPhos (0.094 g, 0.203 mmol, 10 mol%) and Pd_2_(dba)_3_ (0.046 g, 0.050 mmol, 3 mol%) at room temperature. The resulting reaction mixture was irradiated at 150 °C for 30 min in microwave. The reaction mixture was diluted with H_2_O (100 mL) and was extracted with EtOAc (200 mL × 2). The combined organic extracts were washed with brine, dried over anhydrous Na_2_SO_4_ and concentrated under reduced pressure. The crude product was purified by silica gel column chromatography, using MeOH: CH_2_Cl_2_ (gradient elusion 0:1 to 2:8), followed by (C-18) silica gel column chromatography, using MeCN:H_2_O = 0:10 → 3:7 as a gradient, to afford (S)-1-(4-(4-(4-fluorophenyl)-5-(pyridin-4-yl)-1H-imidazol-2-yl)phenyl)-2,4-dimethylpiperazine (**14f**) (0.120 g, 15%) as a pale-yellow solid. Melting point: 264-270 °C; ^1^H NMR (400 MHz, DMSO-*d*_*6*_ + one drop of TFA) δ 9.69 (s, 1H), 8.76 (d, *J* = 5.9 Hz, 2H), 8.03 (d, *J* = 8.7 Hz, 2H), 7.98 (d, *J* = 6.5 Hz, 2H), 7.69 (dd, *J* = 8.4, 5.5 Hz, 2H), 7.44 (t, *J* = 8.8 Hz, 2H), 7.14 (d, *J* = 8.8 Hz, 2H), 4.54 (s, 1H), 3.88 (d, *J* = 12.8 Hz, 1H), 3.54 (d, *J* = 11.7 Hz, 2H), 3.36 – 3.05 (m, 3H), 2.89 (s, 3H), 1.16 (d, *J* = 6.9 Hz, 3H); ^13^C NMR (100 MHz, DMSO-*d*_*6*_ + one drop of TFA) δ 163.1 (^1^*J*_*CF*_ = 247.2 Hz), 149.7, 147.7 (2C), 142.4 (2C), 135.3, 131.8 (^3^*J*_*CF*_=8.6 Hz, 2C),, 128.7, 127.8 (2C), 125.5, 122.3 (2C), 117.3, 116.5 (^2^*J*_*CF*_ = 21.9 Hz, 2C), 114.9 (2C), 56.7, 52.7, 47.6, 43.16, 38.4, 11.8.; ^19^F NMR (400 MHz, DMSO-*d*_*6*_ + one drop of TFA) δ −110.9 Hz; HRMS m/z [M+H]^+^ calcd for C_26_H_27_FN_5_: 428.2245, found 428.2245, LC *t*_R_ = 2.18 min, > 98 % purity.

#### *Condensation:* 2-(4-(Diethylamino)phenyl)-4-(4-fluorophenyl)-5-(pyridin-4-yl)-1*H*-imidazol-1-ol (10II)

To a stirred solution of (E)-1-(4-fluorophenyl)-2-(hydroxyimino)-2-(pyridin-4-yl)ethan-1-one (**10**) (0.700 g, 2.86 mmol) in toluene:AcOH (15:5, 20 mL) were added 4- (diethylamino)benzaldehyde (0.507 g, 2.86 mmol, 1.0 equiv) and NH_4_OAc (1.76 g, 22.95 mmol, 8.0 equiv) portion wise at room temperature and the resulting reaction mixture was stirred at 110 °C for 16h. Reaction mixture was cooled to room temperature, diluted with H_2_O (50 mL) and was extracted with EtOAc (50 mL × 4). The combined organic extracts were dried over anhydrous Na_2_SO_4_ and concentrated under reduced pressure. The crude product was purified by silica gel column chromatography, using MeOH: CH_2_Cl_2_ (gradient elusion 0:1 to 2:8), to afford, 2-(4-(diethylamino)phenyl)-4-(4-fluorophenyl)-5-(pyridin-4-yl)-1H-imidazol-1-ol (**10II**) (0.320 g, 27%) as a brown solid. ^1^H NMR (400 MHz, DMSO-*d*_*6*_) δ δ 11.92 (s, 1H), 8.60 (d, *J* = 6.0 Hz, 2H), 7.91 (d, *J* = 8.8 Hz, 2H), 7.53 – 7.46 (m, 2H), 7.43 (dd, *J* = 4.5, 1.6 Hz, 2H), 7.29 (s, 2H), 7.16 (t, *J* = 8.9 Hz, 2H), 6.81 – 6.64 (m, *J* = 14.4 Hz, 4H), 3.39 (dd, *J* = 14.0, 7.0 Hz, 4H), 1.12 (t, *J* = 7.0 Hz, 6H); MS: [MH]^+^ 402.9.

#### *Dehydroxylation:* N,N-diethyl-4-(4-(4-fluorophenyl)-5-(pyridin-4-yl)-1*H*-imidazol-2-yl)aniline (16)

To a stirred solution of 2-(4-(diethylamino)phenyl)-4-(4-fluorophenyl)-5-(pyridin-4-yl)-1H-imidazol-1-ol (**10II**) (0.300 g, 0.746 mmol)) in DMF (3 mL) was added P(OEt)_3_ (128 μL, 0.746 mmol) drop wise at room temperature and the resulting reaction mixture was stirred at 110 °C for 4h. The reaction mixture was cooled to room temperature, diluted with H_2_O (50 mL) and extracted with EtOAc (50 mL × 3). The combined organic extracts were washed with brine, dried over anhydrous Na_2_SO_4_ and concentrated under reduced pressure. The crude product was purified by silica gel column chromatography, using MeOH: CH_2_Cl_2_ (gradient elusion 0:1 to 2:8), to afford N,N-diethyl-4-(4-(4-fluorophenyl)-5-(pyridin-4-yl)-1H-imidazol-2-yl)aniline (**16**) (0.050 g, 17%) as a light yellow solid. Melting point: 255-270 °C; ^1^H NMR (400 MHz, DMSO-*d*_*6*_ + one drop of TFA) δ δ 8.84 (d, *J* = 6.3 Hz, 2H), 8.08 (d, *J* = 8.3 Hz, 2H), 8.00 (d, *J* = 6.5 Hz, 2H), 7.70 (dd, *J* = 8.6, 5.4 Hz, 2H), 7.44 (t, *J* = 8.8 Hz, 2H), 7.16 (br s, 2H), 3.52 (dd, *J* = 13.8, 6.8 Hz, 4H), 1.10 (t, *J* = 7.0 Hz, 6H); ^13^C NMR (100 MHz, methanol-*d*_*4*_) δ 164.2 (^1^*J*_*CF*_ = 247.2 Hz), 150.6, 150.0 (2C), 149.9, 132.1 (^3^*J*_*CF*_=8.2 Hz, 2C), 128.4 (2C), 123.0 (2C), 117.4, 116.8 (^2^*J*_*CF*_ = 22.0 Hz, 2C), 112.5 (2C), 45.4 (2C), 12.9 (2C).; ^19^F NMR (400 MHz, DMSO-*d*_*6*_ + one drop of TFA) δ −110.4 Hz; HRMS m/z [M+H]^+^ calcd for C_24_H_24_FN_4_: 387.1980, found 387.1997, LC *t*_R_ = 3.26 min, > 98 % purity.

#### Mass spectrometry

Samples were analysed with a UHPLC-qTOF-MS system (1290 UHPLC, Jetstream ESI source, 6540 UHD qTOF-MS, Agilent Technologies, Waldbronn, Karlsruhe, Germany). Data acquisition software was MassHunter Acquisition B.04.00 (Agilent Technologies). Two microliters of the sample solution were injected onto a column (Acquity UPLC BEH C18 column, 2.1×50 mm, 1.7 μM, Waters Corporation, Milford, MA, USA) that was kept at 60 °C. Mobile phases, delivered at 500 µL/min, consisted of water (eluent A) and acetonitrile (eluent B), both containing 0.1 % (v/v) of HCOOH. The following gradient profile was used: 0–9.8 min: 2 → 95% B, 9.8–10.2 min: 95% B, 10.20–10.21 min: 95 → 2% B; 10.21–12.5 min: 2% B. The sample tray was kept at 10 °C during these analyses.

A Jetstream ESI source, operated in positive electrospray ionization (ESI+) mode, used the following conditions: drying gas temperature 325 °C and a flow of 10 L/min, sheath gas temperature 350 °C and a flow of 11 L/min, nebulizer pressure 45 psi, capillary voltage 3500 V, nozzle voltage 1000 V, fragmentor voltage 100 V, and skimmer 45 V. Nitrogen was used as the instrument gas. For data acquisition, a 2 GHz extended dynamic range mode was used with a mass range from m/z 20 to 1600. Data was collected in the centroid mode at an acquisition rate of 1.67 spectra/switch with an abundance threshold of 150. The TOF was calibrated daily and subsequently operated at high accuracy (<2 ppm). Continuous mass axis calibration was performed by monitoring two reference ions from an infusion solution throughout the runs. The reference ions were m/z 121.050873 and m/z 922.009798.

Mass Hunter Qualitative Analysis Navigator (Version B.08.00, Agilent Technologies) was used to analyze the data. Solutions were analyzed at 5 µg/mL or 50 µg/mL based on responsiveness to the ESI mechanism. All observed species were singly charged, as verified by unit m/z separation between mass spectral peaks corresponding to the 12C and 13C12Cc-1 isotope for each elemental composition.

## Acknowledgements

We thank the Academy of Finland PROFI6 program for financial support (CRMA). We thank Biocenter Finland/DDCB for financial support and CSC – IT Center for Science, Finland, for computational resources (TL and AP). In addition, we thank the Finnish Ministry of Education and Culture for allocation of computational resources on the Mahti supercomputer (TL and AP). In addition, we thank Biocenter Finland/Metabolomics for financial support and Erika Pennanen, Miia Reponen and Marko Lehtonen for the collection and processing of the LC-HRMS data.

